# Computational modeling of head direction cells in three-dimensional space: directional encoding and visual cue manipulation

**DOI:** 10.64898/2026.02.02.703434

**Authors:** Yihong Wang, Jianbiao Hu, Shuang Xu, Xuying Xu, Xiaochuan Pan, Rubin Wang

**Affiliations:** Institute for Cognitive Neurodynamics, School of Mathematics, East China University of Science and Technology, Shanghai, China; Center for Intelligent Computing, School of Mathematics, East China University of Science and Technology, Shanghai, China; School of Computer Science and Technology, Hangzhou Dianzi University, Hangzhou, China

**Keywords:** Spatial Navigation, Neurodynamics, Head Direction Cell, Continuous Attractor Network, Three-Dimensional Space

## Abstract

Head direction (HD) cells can fire selectively as a function of the animal’s head azimuth direction and form an internal compass for navigation. They were found in the mammalian limbic system including the dorsal presubiculum and entorhinal cortex. The underlying network updates its directional estimate in a self-organized fashion and can be recalibrated by external sensory cues. Although ring-attractor models were proposed to account for azimuth coding on horizontal planes, they cannot explain conjunctive azimuth-and-pitch tuning observed in HD cells of the presubiculum of flying bats navigating in three-dimensional (3-D) space. Based on the 3-D electrophysiological recordings, we developed a toroidal continuous attractor network that jointly encodes horizontal azimuth and vertical pitch angle of head direction. The model can reproduce the experimentally recorded tuning curves of individual HD cells, and accurately encode the 3-D dynamic head direction of bat by HD cell population. The model also simulate the influence of horizontal visual cue manipulation on the HD system in 3-D space and predicts how horizontal landmark rotation induces a sustained azimuthal offset that persists after the cue is removed, which is comparable to two-dimensional experimental findings. This research clarifies how conjunctive 3-D direction codes are generated and modified by vestibular input, visual information and recurrent connectivity. It also uncovers the computational principles and properties of the brain’s navigation functions in realistic 3-D environments and offers new theoretical reference for future studies.

**Author summary:** A central function of the brain’s navigation system is to track head direction, which in terrestrial mammals is largely confined to a horizontal plane. However, for animals like bats that navigate freely in three-dimensional (3-D) space, the neural code for head direction has been found to be high-dimensional. While ring attractor network models elegantly explain horizontal azimuth coding, how the 3-D head direction code is formed and affected by sensory information remains unknown. Here, we develop a toroidal continuous attractor network model with vestibular and visual modules that explains the jointly encoding of azimuth and pitch angles, and the effect of visual cue. We introduce the toroidal topology to the continuous attractor network, and the high-dimensional angular velocity signal from vestibular system and visual input are used to update the population dynamics of the network. Our model is constrained by and reproduces 3-D electrophysiological data from flying bats, captures the unique tuning curves of individual head direction cells, and further achieves the accurate encoding of dynamic 3-D head movements by population activity. Evidences in the rodent have suggested visual cue manipulation recalibrates head direction coding horizontally. Our model further successfully simulates the influence of visual cue rotation in 3-D space, predicting a persistent, global offset in the azimuth encoding—an effect that endures after cue removal and aligns with 2-D rodent experimental phenomena. This work provides a mechanistic, network-level explanation for 3-D head direction encoding and reveals how multimodal cues are integrated to form the internal compass in a volumetric world.

## Introduction

Navigation in natural environments is crucial for animals’ survival and depends critically on the encoding of multimodal spatial information by distinct neuronal populations. Principal neurons in mammalian hippocampal system are believed to encode location, time and memory information (1), forming a ‘cognitive map’ to support flexible navigation (2). Since navigation occurs in the approximately Euclidean space, which has not only location but also direction, an internal compass is needed to maintain the sense of direction while navigating. Head direction (HD) cells were initially identified in the dorsal part of hippocampal presubiculum of freely moving rats that exhibited robust responses to specific head directions (3, 4), which function together as a neural compass. These cells have since been found in other brain regions, including the anterior thalamic nuclei (5, 6) and the deep medial entorhinal cortex (7) of mammals. Similar signals representing facing direction have also been found in human retrosplenial complex and superior parietal lobule (8). An HD cell fires most intensively when the animal’s head points toward a specific direction, defined as the cell’s preferred direction. The firing rate of a HD cell as a function of head direction usually forms a unimodal tuning curve (5, 6). Other than the static directional tuning features, recent discoveries of HD cells in the anteroventral thalamic nucleus of rodents have further revealed theta-modulated phase precession (9). External sensory cues can strongly influence the directional tuning of HD cells. For example, rotating distal visual landmarks in the horizontal plane shifts the preferred directions of these cells. In contrast, the animal’s intrinsic physical characteristics (e.g., body height and weight), ongoing behavior, and spatial location within the environment rarely impact HD cell activity (3, 5).

The physical world is inherently three-dimensional (3-D), so the 3-D orientation encoding is required by the animal for successful spatial navigation. How the brain represents three-dimensional direction is a fundamental question in neuroscience. Early studies identified directionally tuned neuronal activity in rodents navigating on different surfaces in 3-D space (10-12). Subsequent experiments revealed neurons sensitive to head pitch (or tilt) in the lateral mammillary nucleus (13) and anterior thalamus (14) of rats, the anterior thalamus and retrosplenial cortex of mice (15), and the anterior thalamus of macaques (16). While these animals typically require additional physical structures to navigate in 3-D space, bats—which navigate freely and actively in three dimensions—provide a natural model for studying 3-D spatial cognition (17, 18). Recordings from the presubiculum of bats have identified HD cells both in natural environments (19) and in laboratory settings (20). Evident 3-D tuning properties of HD cells have been revealed for freely flying or crawling bats (20). Using concepts from rigid body mechanics, 3-D head orientation in bats can be decomposed into three Euler angles: azimuth (yaw), pitch, and roll (21). Unlike rodents such as rats, flying mammals like bats exhibit more frequent and unconstrained head movements, particularly in pitch, which can vary freely through a full 360° range. The tuning properties of bat HD cells also differ from those of rodents. For instance, HD cells in bats maintain directional selectivity even during inverted postures (20), whereas those in rats largely lose their directional tuning under similar conditions (13, 14). Moreover, the bat presubiculum contains not only neurons tuned selectively to individual Euler angles but also cells that conjunctively encode pairs of angles—or even all three Euler angles simultaneously (20). Notably, some neurons tuned to a specific azimuth during upright (ventral-down) posture shift their preference 180° to the opposite allocentric direction when the bat is inverted (ventral-up). These findings pose challenges for defining neuronal preferred directions using spherical topology (22-26), as two major issues arise: (1) the preferred azimuth of such neurons exhibits a 180° shift between inverted and upright postures, and (2) singularities in azimuth tuning emerge at the north and south poles (azimuth angle becomes ill-defined) of the spherical coordinate system.

When visual cues are available, the HD system typically anchors to landmarks. Rotations of visual cues on the horizontal plane induce corresponding shifts in the internal directional representation of the HD system (6, 27-29). In rats, such rotations introduce systematic decoding errors that match the cue displacement and persist after the cue is removed (30). This finding indicates that when salient visual cue conflicts with self-motion signal integration, rats rely more confidently on visual information, leading to a reset of their internal directional representation. However, whether similar mechanisms govern 3-D directional encoding in bats is unknown.

Here we address three unanswered questions. (i) How does the population of HD cells in bats integrate self-motion perception and environmental landmarks to jointly encode the three Euler angles of 3-D head orientation? (ii) How are the 180° azimuth reversal at inversion and the spherical-coordinate singularity resolved at the neuronal level? (iii) Can landmark manipulation elicit systematic 3-D directional errors similar to those observed in horizontal-plane experiments?

To explain the activity of spatial neurons, many classic models have been developed. Continuous attractor dynamics have been proven to be a promising and plausible mechanism underlying the encoding for many continuous physical variables by animal’s neural system (31). Its theoretical framework has been used in many spatial cognition modeling such as modeling place cell (32, 33), grid cell (34-36) and head direction cell (22, 37, 38). Experimental data recorded across species (39-41) and emerging analytical methods (42, 43) increasingly support the continuous attractor architecture as a reliable theoretical model framework for navigation circuits. And its encoding ability for location and direction have been used for brain-inspired robot navigation system (44). But how the 3-D head direction encoding of bats can be incorporated to this theory remains unknown. To address how the 3-D neural compass operate in the bat’s brain, we developed a continuous attractor network model of 3-D HD cells, focusing on neurons in the bat presubiculum that conjunctively encode azimuth and pitch. Due to the fact that bats seldom perform roll movements under natural conditions, and HD cells tuned to roll angle are relatively scarce (20, 45), we chose to neglect the roll encoding to reduce the model’s complexity without losing too much fidelity (in the recorded neurons, 12.3% have roll sensitivity and 4.9% are pure roll selective (20)). The model consists of three modules: a self-motion input (vestibular signals) module, a visual cue input module, and a HD cell network modeling the bat’s presubiculum. The HD layer is constructed as a two-dimensional neural sheet with periodic boundary conditions along both axes, forming a toroidal topology that can be viewed as the Cartesian product of two ring attractor networks.

The model can accurately integrate angular velocity in both azimuth and pitch in the absence of visual input, maintaining a stable internal representation of these two Euler angles. Individual neurons in the HD cell network exhibit conjunctive tuning curves (surfaces) to both azimuth and pitch. When distal landmark (visual cue) is introduced and systematically rotated in the azimuthal direction (note: pitch perception is primarily anchored to gravity (15, 16, 46, 47) and cannot be easily manipulated, so only azimuthal visual cue was rotated in the model), the decoded population activity of HD cells shifts accordingly, similar to experimental observations in rats (30). Concurrently, the preferred azimuth of individual HD cell also rotates coherently with the visual cue. Our model indicates that toroidal attractor dynamics can explain the emergence of conjunctive azimuth-pitch tuning in 3-D space observed in bats, effectively resolves the azimuth shift and singularity problems. It further predicts that visual cue manipulations induce similar systematic errors in azimuth encoding while the toroidal neural manifold may retain stable. These findings offer novel insights into the computational principles underlying 3-D directional encoding in natural environments.

## Model and method

The model is motivated by recent electrophysiological evidence that presubiculum HD cells in freely flying bats conjunctively encode horizontal azimuth and vertical pitch (20). In this model, we define the bat’s head direction using three Euler angles—azimuth, pitch, and roll, as illustrated in Fig 1a. The architecture of the model is depicted in Fig 1b. Its central component is a network of HD cells in the dorsal presubiculum of bat, arranged into a two-dimensional neural sheet (Fig 1c), where the horizontal and vertical axes represent preferred directions of azimuth and pitch respectively (note that this specific assignment is not unique, in fact, the two dimensions exhibit structural symmetry). Each HD cell on the neural sheet receives three types of input: 1) input by recurrent connections from other HD cells within the dorsal presubiculum network, 2) head angular velocity information from the vestibular system driving the angular path integration, and 3) visual anchoring cues provided by visual system.

**Fig 1.**
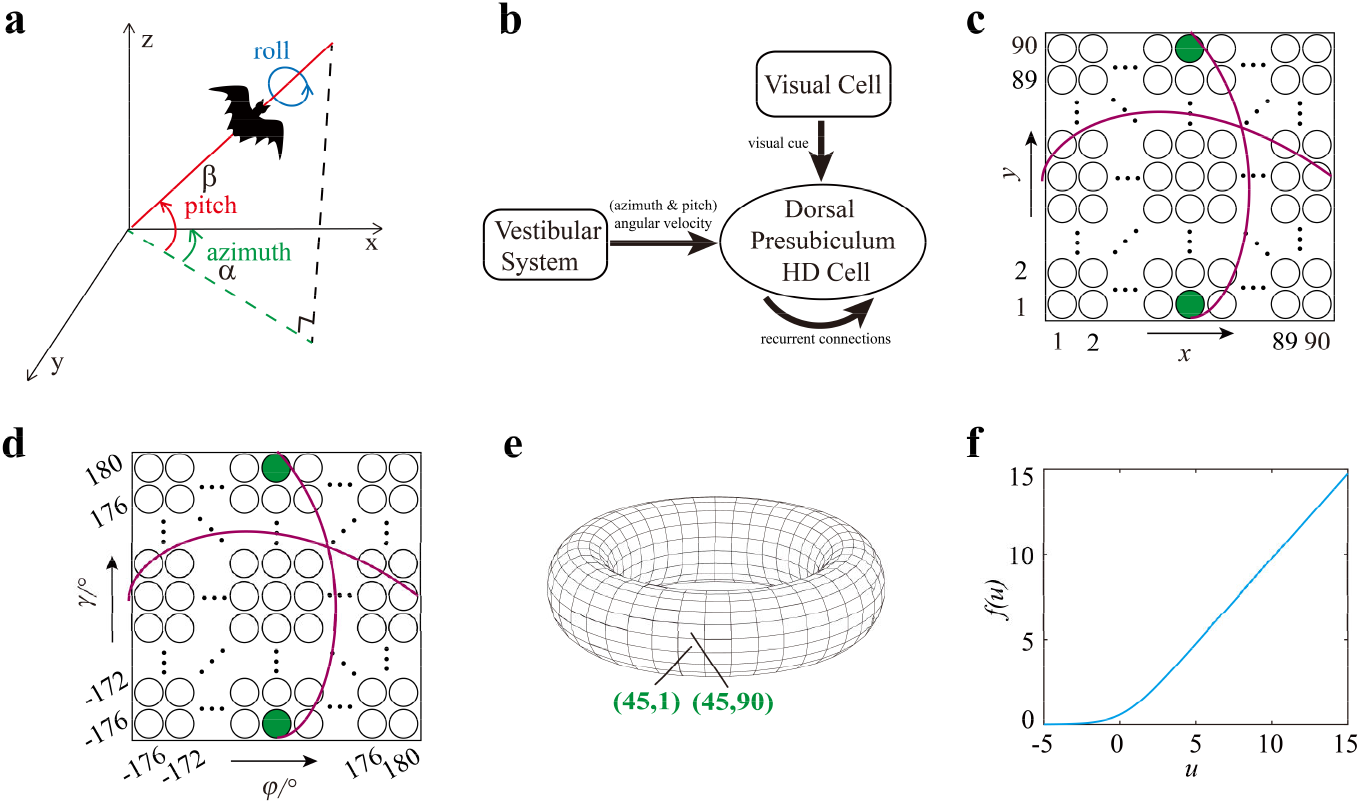
The framework and structure of the HD cell model in 3-D space. (a) Definitions of Euler angles of a bat in three-dimensional space. When the bat’s head rotates counterclockwise around A-P axis along the blue circle, the roll angle (blue arrow) increases. When the head rotates counterclockwise around the z-axis, the azimuth angle (green arrow) increases. When the head tilts above the horizontal plane, the pitch angle (red arrow) increases. (b) Model architecture: The HD cell population forms a recurrent network. Each HD cell receives (azimuth and pitch) angular velocity signals of head rotation from the vestibular system and may also receive visual input depending on experimental condition. (c) 8100 HD cells are arranged on a 90×90 sheet. The x and y axes correspond to azimuth and pitch indices, respectively, ranging from 1 to 90 (left to right and bottom to top). HD cells at opposite ends of the purple lines (along either axis) encode adjacent directions. (d) Preferred direction tuning of HD cells arranged on the 90×90 sheet. Each cell’s preferred azimuth *φ* and pitch *γ* are determined by its grid position. *φ* increases from -176° to 180° along the horizontal axis (left to right), and *γ* increases from - 176° to 180° along the vertical axis (bottom to top), with a 4° increment between adjacent cells. (e) Periodic connectivity in the HD network topology. As shown in panel (c), the two green-labeled HD cells reside at opposite boundaries of the vertical (pitch) axis. Under periodic boundary conditions, these cells become adjacent on the toroidal manifold. (f) Neuronal activation function.

The network contains *N* = 8100 HD cells, arranged as a 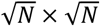 two-dimensional neural sheet. Each HD cell *i* (*i* = 1, 2, …, *N*) is assigned a 2-D coordinate (*x*_*i*_, *y*_*i*_) within the sheet, from (1, 1) to (90, 90), as shown in Fig 1c. Each neuron *i* encodes a preconfigured preferred direction ***ϑ***_*i*_ (which will be anchored to the external physical orientation through visual information), defined as:

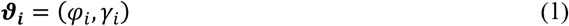

Where *φ*_*i*_ and *γ*_*i*_ denote the preferred azimuth and preferred pitch of neuron *i* respectively. The values of *φ*_*i*_ and *γ*_*i*_ are discretized as −176°, −172°, −168°, …, 180° (Fig 1d). To implement periodic boundary conditions, the neurons on the left and right edges as well as that on the top and bottom edges of the neuron sheet are connected, forming a toroidal connection topology. For instance, cells at positions (45, 1) and (45, 90) (highlighted in green in Fig 1c and 1d) become adjacent on the torus (Fig 1e, black dots). The firing rate *r*_*i*_ of neuron *i* follows the dynamics (34):

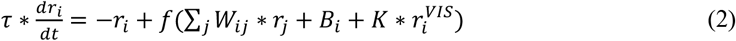

Where *τ* is the time constant (6ms in this model), *W*_*ij*_ is the synaptic weight from neuron *j* to neuron *i, B*_*i*_ represents vestibular angular velocity inputs, 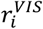 denotes visual inputs, and *K* is a gating variable dependent on visual conditions in experiments (exists in sight or not). The activation function *f*(*u*) is defined as:

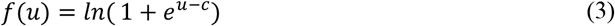

with *c* = 0.25. As shown in Fig 1f,when input *u* ≤ 0,*f*(*u*) ≈ 0, and *f*(*u*) ≈ *u* for positive *u*.

To enable the HD network to integrate vestibular inputs, each HD cell *i* is assigned a preferred rotation direction *δ*_*i*_ ∈ {*PI, AI, PD, AD*} (Fig 2a). Each adjacent 2×2 block of HD cells on the neural grid uniformly samples all four directions *PI, AI, PD, AD* (Fig 2b). *δ*_*i*_ determines: 1) the direction of the output weight center offset for HD cell *i* (relative to the center-symmetrical local excitation and long-range inhibition weights (Fig 2c), and 2) the differential response of HD cell *i* to angular velocity input which signaling changes in head direction. Here, *AI* and *AD* correspond to increase and decrease in azimuth, respectively, while *PI* and *PD* correspond to increase and decrease in pitch, respectively (Fig 2c). The synaptic weight distribution approximates a classical center-excitation, surround-inhibition profile (a 1-D example without preferred rotation offset is shown in Fig 2d), ensuring that the activity bump solution of the HD cell population remains unimodal (48). The connection weight *W*_*ij*_ from HD cell *j* to HD cell *i* satisfies:

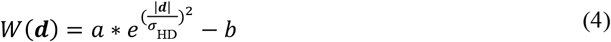

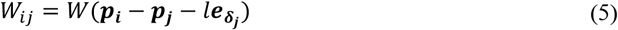

where *b* determines the strength of inhibitory connections, and *σ*_*HD*_ determines the width of the excitatory connection weight distribution. We set *a* = 0.1205 and *b* = 0.112, allowing the HD cell population to maintain stable collective activity. *σ*_*HD*_ = 40 ensures that the range of excitatory connections is approximately 120°, which is biologically plausible. For a presynaptic HD cell *j* at position ***p*** _***j***_ = (*x*_*j*_, *y*_*j*_) on the neural sheet (Fig 1c), the center of its output weight distribution is not centered at ***p***_***j***_, but is offset slightly along the preferred rotation direction *δ*_*j*_ of cell *j*. The new distribution center is at position 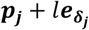, where 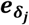 is the unit vector in direction *δ*_*j*_, and *l* = 2 is the offset length, i.e., the center of the output weight distribution is shifted by two grid units along direction *δ*_*j*_ (Fig 2e).

**Fig 2.**
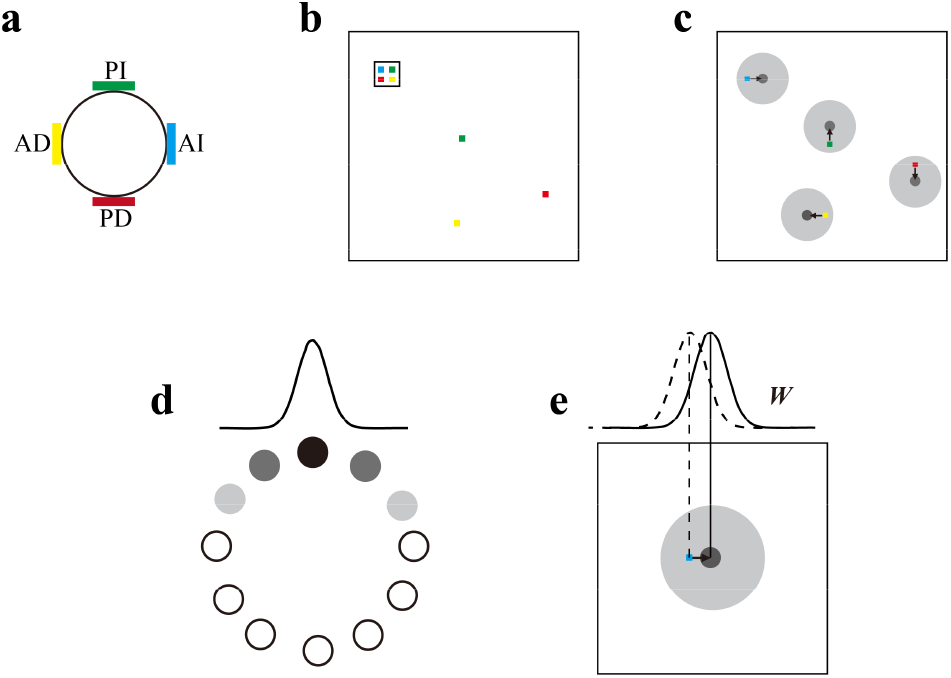
Distribution of connection weights of the HD cells’ network. (a) Offset directions for the output weight distribution centers of HD cells. Blue, yellow, green, and red represent azimuth increase (AI), azimuth decrease (AD), pitch increase (PI), and pitch decrease (AD) rotation directions, respectively (same color code is used in subsequent panels). (b) For the four HD cells within the black square in the upper left of the neural sheet, their weight distribution centers are offset according to the rules in (a). The colored points outside the upper left square show examples of offset directions for other HD cells. (c) Output weight distributions for four example HD cells offsets differently. The darker shade represents the excitatory weight while lighter shade represents inhibitory weight. (d) Schematic of the connection weight distribution between HD cells along one dimension without an offset. And the black fill indicates this particular neuron is firing maximally. Activity decreases as the color lightens. The connection weight profile between HD cells follows the Gaussian-like distribution. (e) Output weight distribution of a HD cell offset to AI direction. The center of its output weight distribution is shifted two units to the azimuth increase direction (dashed line: before shifting; solid line: after shifting).

The angular velocity signal *B*_*i*_ which reflects the head rotation in 3-D space, received by HD cell *i* on the neural sheet is given by:

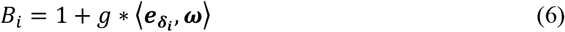

where ***ω*** is the 2-D vector composed of the angular velocities of azimuth and pitch angle; *g* = 4.83 is the gain coefficient determining the strength of angular velocity input to the HD system; and 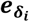 is the unit vector along the preferred offset direction *δ*_*i*_ of HD cell *i* (When *l* = 0 and *g* = 0, the HD cell population could form a stable and static activity bump at any location on the neural sheet). As described earlier, the preferred rotation direction *δ*_*i*_ of HD cell *i* determines both the center of its output weight distribution and the effect of the angular velocity input it receives. This mechanism enables the HD network to integrate angular velocities in 3-D space through updates of the activity bump, thereby encoding the 3-D head direction. This process is similar to path integration in grid cells on a 2-D plane, but the neural sheet does not allow multiple bumps and the resulting spatial periodicity (34).

The model further incorporates a layer of visual (VIS) cells encoding the position of a visual stimulus (mimics a distal light spot or light bar on the horizontal plane in animal experiment) within the field of view. The encoding of vertical pitch angle relies more on gravity perception to anchor (6, 15, 22, 23), while the encoding of horizontal azimuth angle can be influenced by visual stimuli. Therefore, when studying the effect of visual cue manipulation on 3-D head direction encoding, the cue is manipulated only in the horizontal direction. At time *t*, the animal’s actual 3-D head direction is ***θ***_***HD***_ = (*α, β*) (see Fig 1a), and the allocentric direction of the visual cue is ***θ*_*VIS*_** = (*α*_*VIS*_, *β*_*VIS*_), where the horizontal azimuth is *α*_*VIS*_ and the vertical pitch *β*_*VIS*_ is always 0° (assuming the visual stimulus remains on the horizontal plane). Thus, the angular difference ***θ*_*VAR*_** between the animal’s actual head direction ***θ*_*HD*_** and the visual cue direction ***θ*_*VIS*_** at time *t* is:

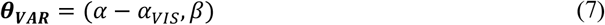

This variable describes the position of the visual cue within the animal’s view field. The VIS cell layer also consists of *N* = 8100 cells arranged on a 90×90 sheet, consistent with the HD cell layer. For any VIS cell *i*, its position on the VIS neural sheet is similarly denoted as (*x*_*i*_, *y*_*i*_) (Fig 3a). Each VIS cell *i* is assigned a preferred view field direction ***Φ***_*i*_ similar to the HD cell *i* (Fig 3b):

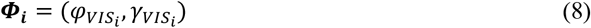

**Fig 3.**
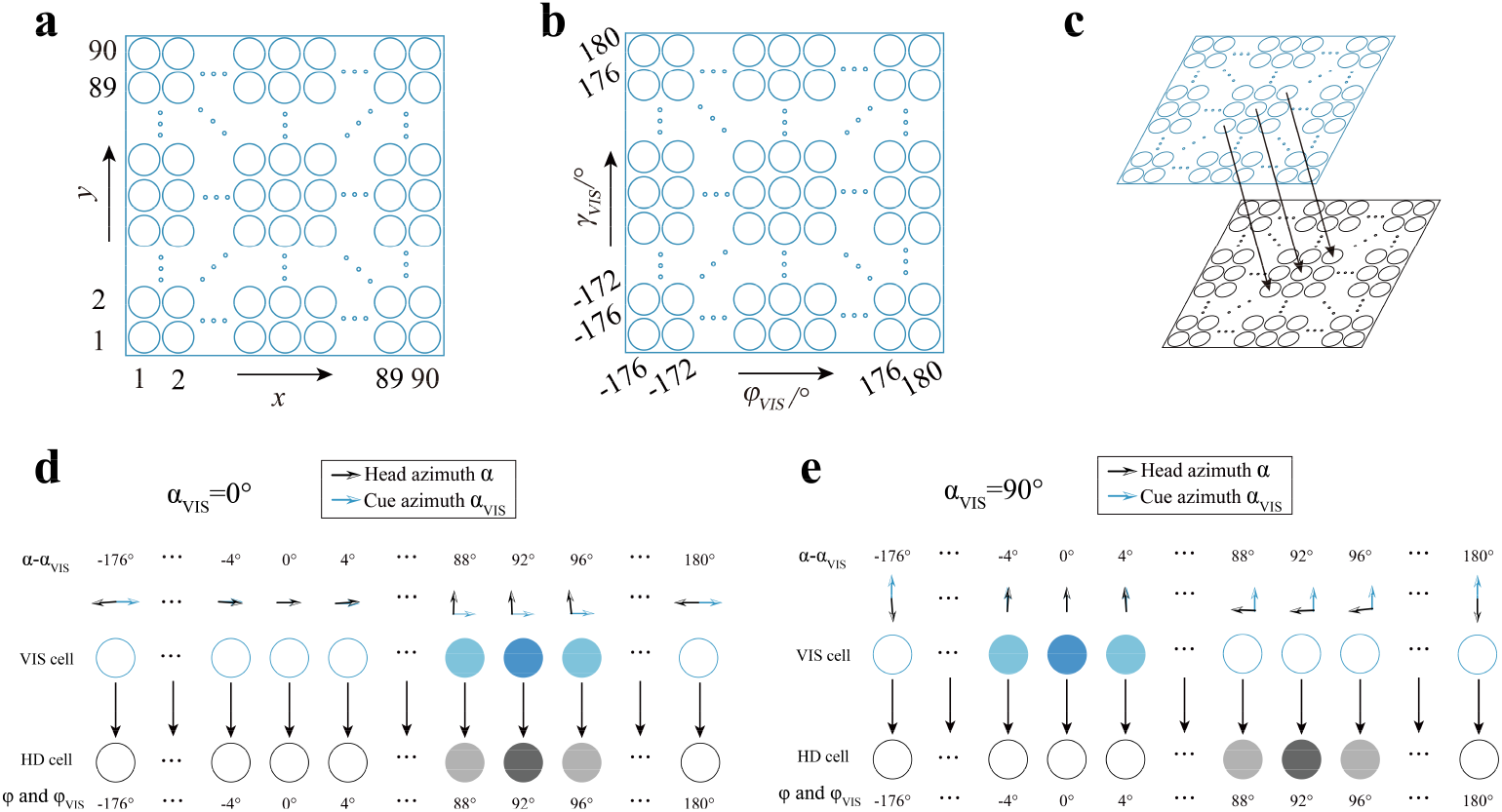
VIS cells encode visual information and form one-to-one connection with HD cells. Only the effect on the horizontal azimuth encoding by the visual information is considered. The virtual visual cue is fixed at a specific azimuth angle. (a) 8100 VIS cells are arranged on a 90×90 sheet. The x and y axes represent the azimuth and pitch indices, respectively, increasing from 1 to 90 from left to right and bottom to top. (b) Each VIS cell is assigned a preferred azimuth and pitch, from -176° to 180°, respectively, same as the HD cells. The incremental step size is 4° for both dimensions. (c) The blue neural sheet represents the VIS cell layer, and the black one represents the HD cell layer. Black arrows illustrate examples of one-to-one connections between VIS and HD cells. (d) and (e) show exampling activity profiles of VIS and HD cells under the same initialization condition of the visual cue direction ***θ*_*VIS*_** = (0^°^, 0^°^), with current head azimuth *α* ≈ 90° and current cue azimuth *α*_*VIS*_ equal to 0° (d) or 90° (e). Blue circles denote VIS cells, and black circles denote HD cells. Darker color indicates higher firing activity. (d) When the visual cue azimuth *α*_*VIS*_ is approximately 0°, HD cells with preferred directions near 90° exhibit the strongest activity, while VIS cells with a preferred azimuth difference of about 90° show the highest response. (e) When *α*_*VIS*_ is 90°, HD cells with preferred directions near 90° still show the strongest activity according to angular path integration, but VIS cells with a preferred azimuth difference of 0° now exhibit the highest response, which provide strong input to HD cells with preferred azimuth of 0° instead of 90°. In (d) and (e), the angle between the two arrows in the second row (representing the azimuth difference between the visual cue and head direction) illustrates the condition under which the corresponding VIS cells respond maximally. The blue and black lines indicate the azimuth angles of the visual cue and head direction, respectively, while their pitch angles remain identical.

As shown in Fig 3c, the VIS cell layer and the HD cell layer are connected in a one-to-one manner according to their preferred directions. The firing rate 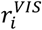 of a VIS cell is determined by the directional relationship between the position of the visual cue in the view field and its preferred view field direction ***Φ***_*i*_, effectively encoding the difference between the current head direction and the visual cue direction (egocentric direction of the visual cue). Specifically, different VIS cells exhibit peak responses when the visual cue appears at different positions in the visual field. Thus, 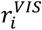 is given by:

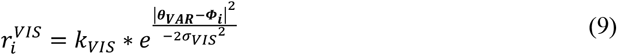

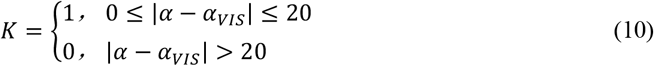

where *k*_*VIS*_ is the peak firing rate of VIS cells, describing the visual input strength and *σ*_*VIS*_ determines the tuning width. Initially, with ***θ*_*VIS*_** = (0^°^, 0^°^), we set *k*_*VIS*_ = 0.3 and *σ*_*VIS*_ = 4 to initialize the HD system, obtaining a stable activity state of the HD cell population. This process can be regarded as binding the internal directional representation of the HD cell population to external environmental information. Under these initial conditions, when the bat’s head direction has an azimuth of approximately 90°, VIS cells with preferred horizontal angular differences near 90° exhibit strong activity, and HD cells with preferred azimuths around 90° also show the strongest responses. Under these conditions, the visual cue and the internal representation remain consistent (Fig 3d). However, when the bat’s head direction azimuth remains at 90° but the visual cue azimuth is shifted from 0° to 90°, a conflict arises between the visual input and the internal representation (Fig 3e). In both examples, the pitch angle of the bat’s head direction and the visual cue remains unchanged for simplicity.

Once the model simulation is finished (with coverage of 360°×360° angular space), the firing rate time series for each HD cell can be obtained. This allows us to determine each HD cell’s empirically preferred direction vector (azimuth and pitch) statistically. For each HD cell *i*, based on its firing rate *r*_*i*_(*t*) and the animal’s actual 3-D head direction ***θ*_*HD*_**(*t*) = (*α*(*t*), *β*(*t*)) at time *t*, its simulation-derived preferred direction vector 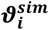 can be computed by taking the firing-rate-weighted average of ***θ*_*HD*_**(*t*) across all time:

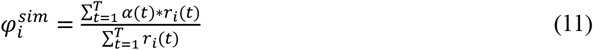

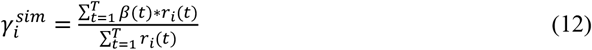

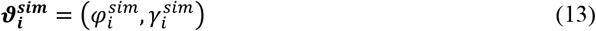

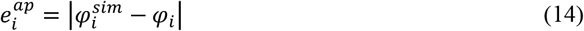

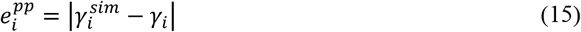

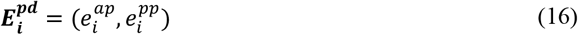

where *φ*_*i*_ and *γ*_*i*_ are the initially assigned preferred azimuth and pitch for HD cell *i* based on its position on the neural sheet; 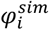 and 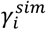 are the preferred azimuth and pitch calculated from simulation data, respectively. 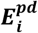 represents the preferred direction error, serving as a metric for the model’s encoding precision of the bat’s 3-D head direction in azimuth and pitch. Based on the population response of HD cells, a decoded head direction 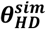 in 3-D space can be obtained using population vector decoding and compared with the true direction ***θ*_*HD*_** at any time *t*:

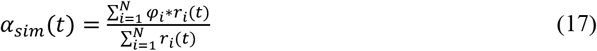

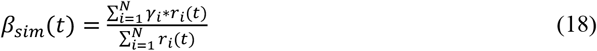

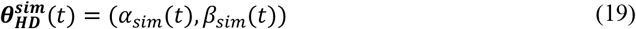

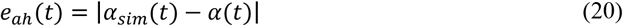

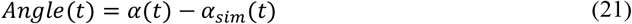

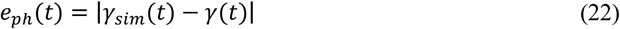

where (*α*_*sim*_(*t*), *β*_*sim*_(*t*)) denotes the head direction vector in 3-D space decoded using the population vector. *e*_*ah*_(*t*) and *e*_*ph*_(*t*) represent the error of azimuth and pitch decoding, respectively. In experiments with rotated visual cues, a persistent error may emerge in azimuth encoding and *angle*(*t*) is defined to quantify this azimuth representation shift error (algebraic value). The key variables and parameters of the model are summarized in Table 1.

**Table 1.**
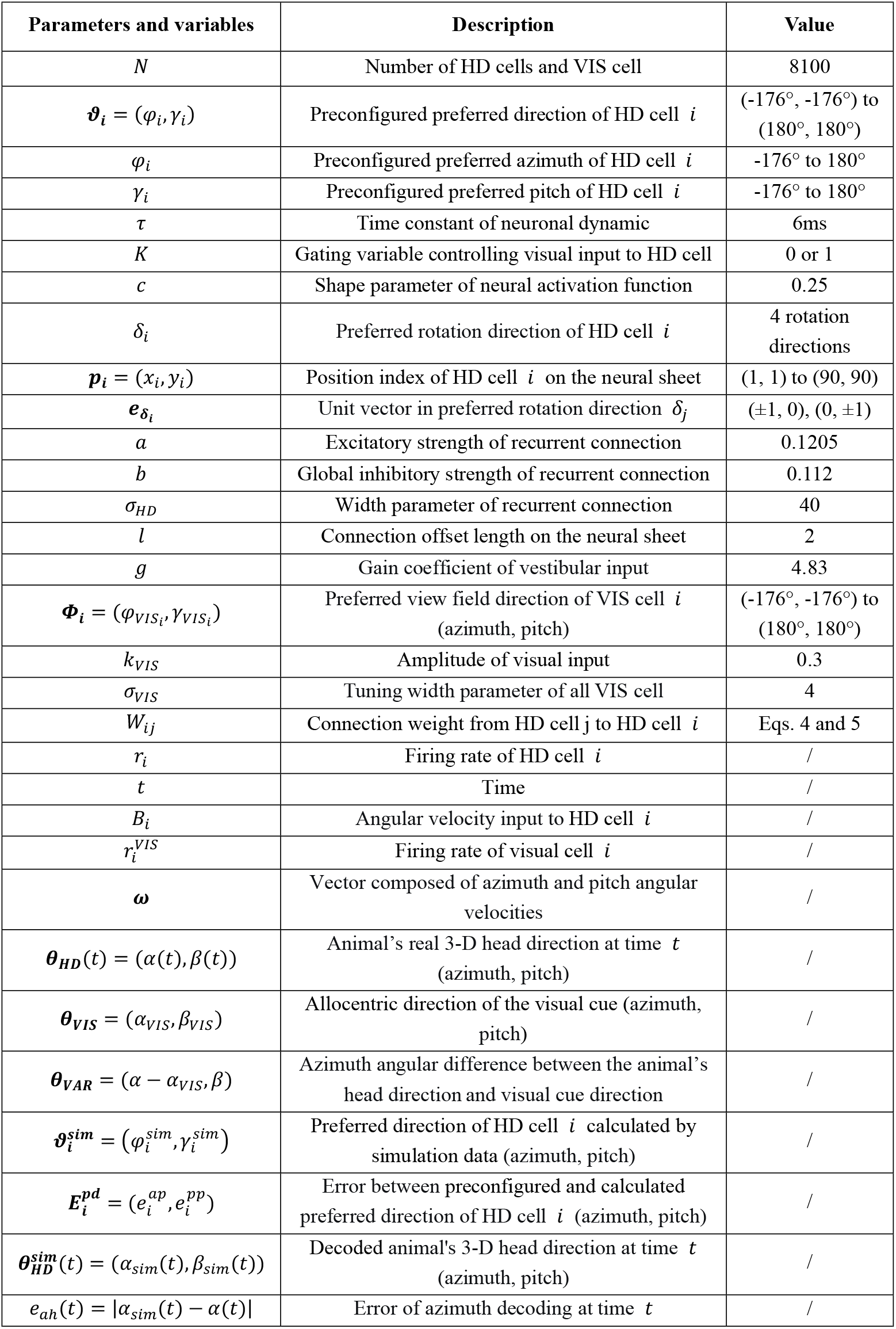
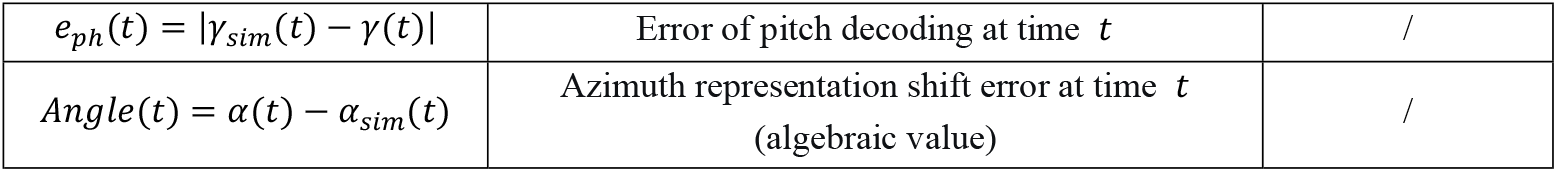
Model variables and parameters.

## Results

### Accurate encoding of direction in 3-D space by HD cell population

The virtual visual cue was positioned at a horizontal azimuth of 0°. The bat’s initial head direction was set to (0^°^, 0^°^). At approximately 400 ms, the HD cell population formed a stable single activity bump at the center of the neural sheet (Fig 4a). The static activity bump at 1000ms was also shown on the neural sheet (Fig 4b) and on the two-dimensional toroidal topology (Fig 4c) after applying periodic boundary condition. Then the HD cells were classified into Fire and Non-Fire groups using a firing rate threshold of 1 (this threshold was also used for all subsequent simulations). Approximately 293 HD cells exhibited significant firing, with a peak firing rate of approximately 6.279 (Fig 4d).

**Fig 4.**
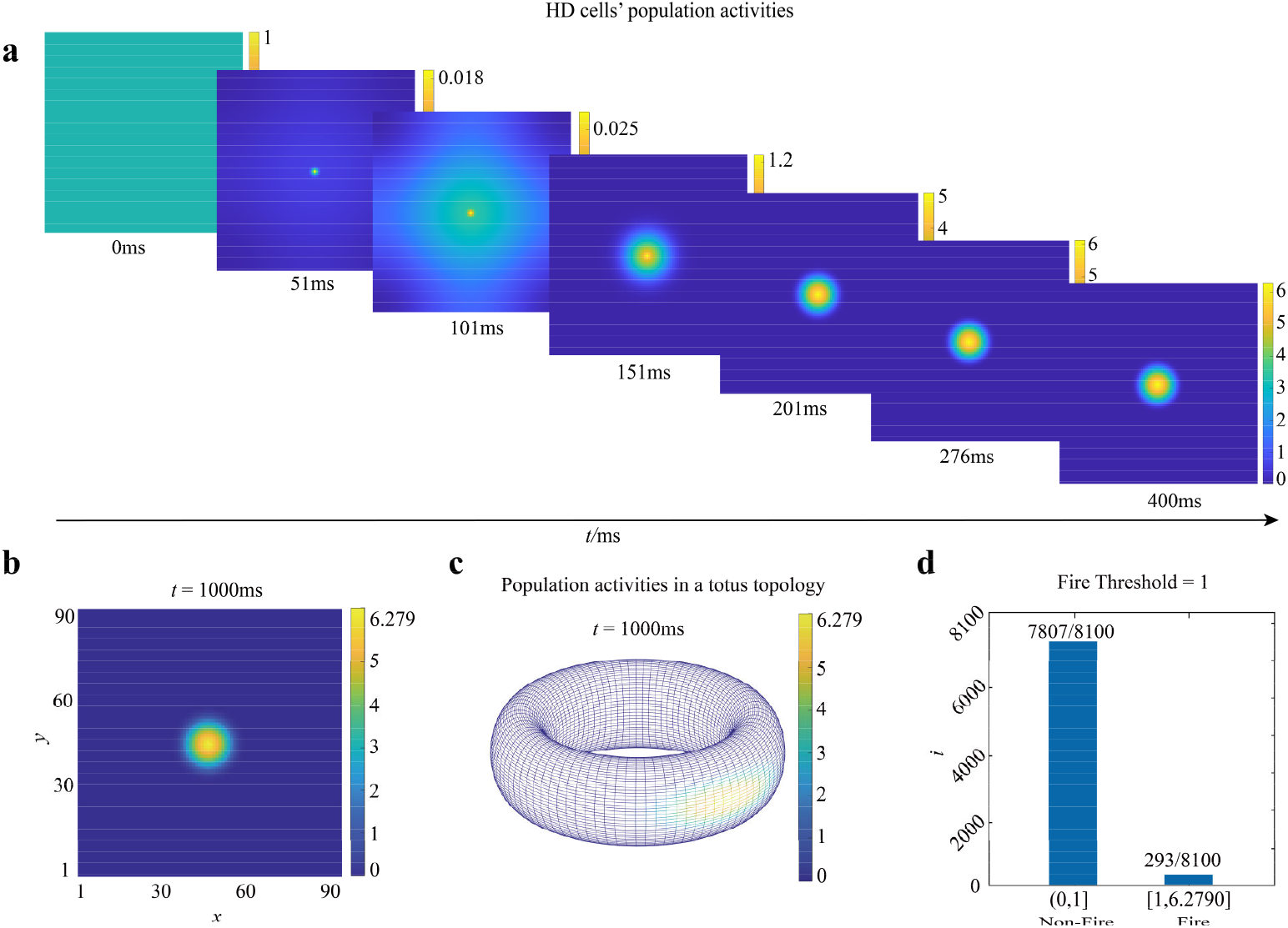
After initialization, single stable activity bump was formed on HD neurons sheet. (a) The evolution of HD cell population activity during 0 to 400 ms. A stable unimodal activity bump gradually formed at the center of the neural sheet. The color bar on the right of the neural sheet indicated the activity intensity. (b) Following initialization of the HD system, the population activity became stable by 1000 ms, forming a static unimodal activity bump precisely at the center of the neural sheet. The maximum firing rate reached approximately 6.279. (c) The neural sheet depicted in panel (b) represented as a toroidal topology. (d) After the HD cell population reached steady state, the firing activity intensity was quantified. Using a firing threshold of 1, cells with activity below this value were classified as Non-Fire, while those above it were considered Fire. Based on this criterion, the activity of all 8100 HD cells was divided into two groups: 293 active HD cells and 7807 inactive ones (activity intensity below threshold).

We first simulated bat flying under two simple conditions, generating motion trajectories in 3-D space containing only changes in head azimuth *α* (with pitch *β* ≡0^°^) and only changes in head pitch *β* (with azimuth *α* ≡0^°^). By inputting the angular velocity vector ***ω*** of head rotation into the model without visual input (only present during initialization as in Fig 4), the HD cell population consistently encoded head direction accurately (Figs 5 and 6).

**Fig 5.**
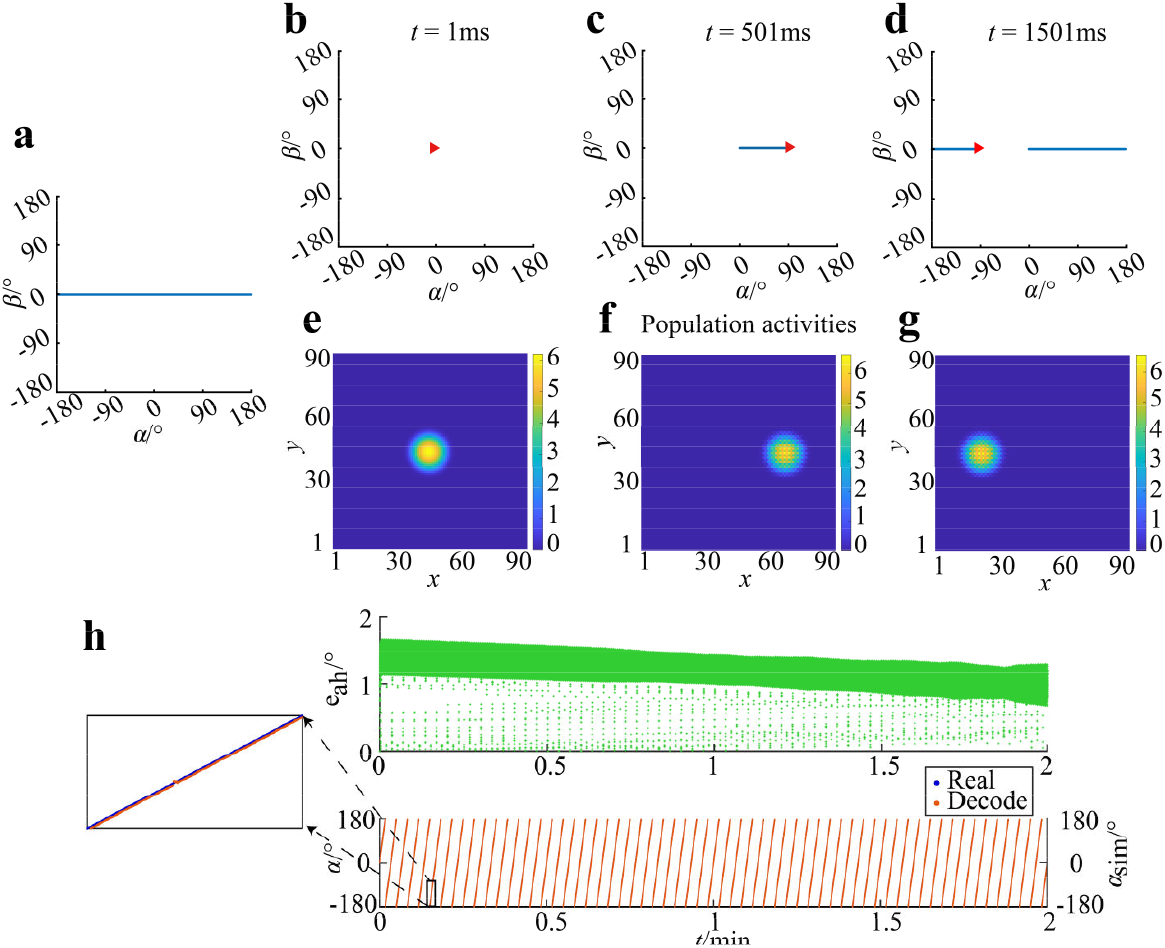
Population dynamics of HD cells accurately tracked changes in head orientation of bats in the horizontal direction. (a) Sequence of bat head direction changes in azimuth within a 2-D subspace (*α*: azimuth, *β*: pitch). Azimuth changes from 0° to 360°, while pitch remains constant at 0°. (b-d) Activity in the 2-D subspace with varying azimuth at 1 ms, 501 ms, and 1501 ms. Red triangles indicate the head direction and bule lines indicate angular trajectory in Euler angular coordinates at the corresponding time. (e-g) Population activity of HD cells on the neural sheet at 1 ms, 501 ms, and 1501 ms. The activity bump moves consistently with the head direction. (h) Decoded azimuth from HD population activity and decoding error *e*_*ah*_. Top: decoding error *e*_*ah*_ (difference between decoded and true azimuth) remains within 2°. Bottom: decoded azimuth *α*_*sim*_ (orange) closely matches true azimuth *α* (blue). The trajectories overlap almost perfectly (blue points are mostly covered by orange points). The inset panel shows a magnified view of the region marked by the black square.

**Fig 6.**
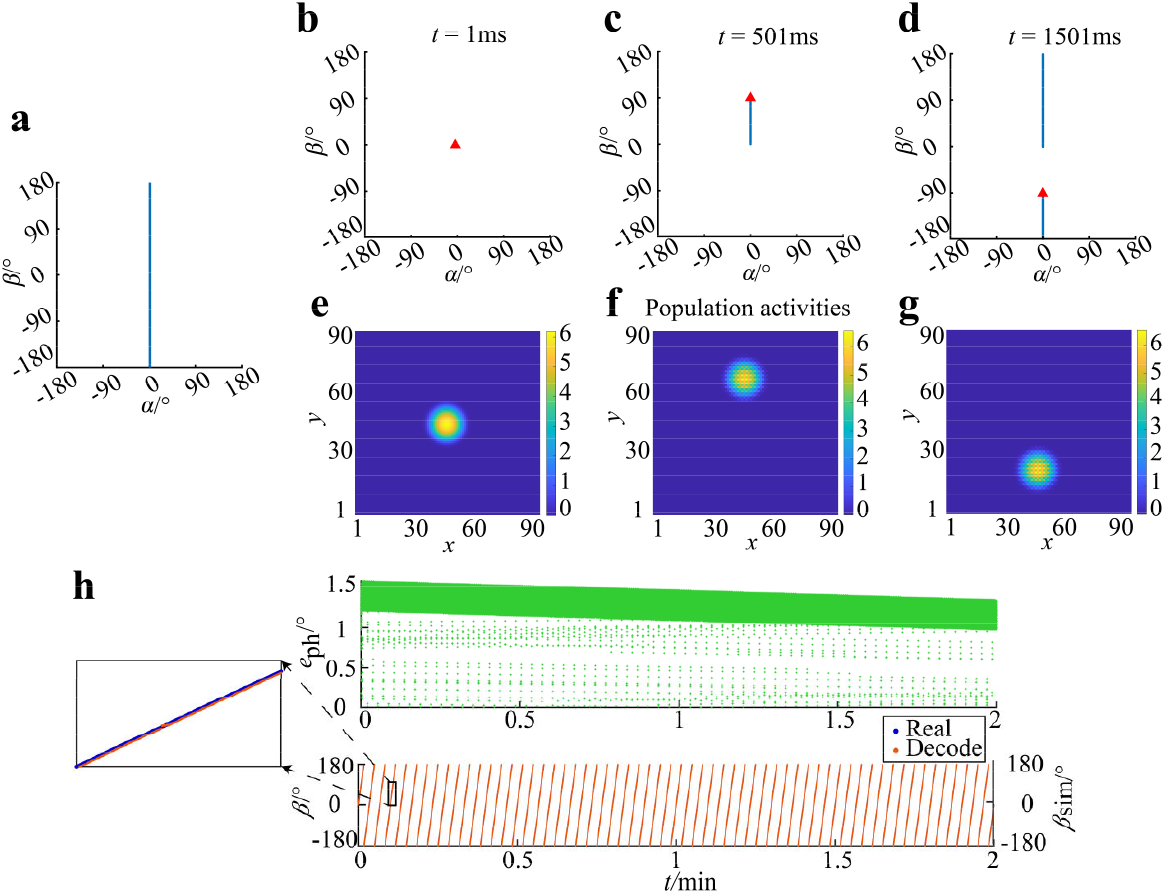
Population dynamics of HD cells accurately tracked changes in head orientation of bats in the vertical direction. All figure labels, axes, and conventions in this figure share the same meanings as in Fig 5, In this case, the azimuth remains constant while the pitch varies from 0° to 360°. The pitch angle was decoded from HD population activity and then compared with true pitch angle.

With pitch fixed at *β* ≡0^°^ and azimuth *α* changing at a constant angular velocity of 0.18°/ms for 2 minutes (Fig 5a), i.e., ***ω*** = (0.18,0)^°^/*ms*, we recorded the position of the activity bump on the neural sheet. The bump movement on the neural sheet matched perfectly the changes in head azimuth *α* in Euler angle coordinates (Figs 5b-g). Using Eq 17 and 18 to decode the population activity, we compared the internally represented head direction 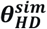 with the true head direction ***θ*_*HD*_** and computed the decoding error *E*_*HD*_. Throughout the 2-minute simulation, the azimuth decoding error remained within 2° (Fig 5h, top), demonstrating accurate encoding of head azimuth by the HD system (Fig 5h, bottom).

Similarly, with azimuth fixed at 0° and pitch changing at a constant angular velocity of 0.18°/ms (Fig 6a), the activity bump also accurately tracked changes in pitch (Figs 6b-g) and the pitch decoding error remained within 1.5° during the 2-minute simulation (Fig 6).

Subsequently, we simulated a longer, more realistic flight trajectory of a bat in 3-D space by generating 40-minute of biologically plausible random motion. The angular velocity ***ω*** of head changes at each time point was input to the model (without visual cue). The results demonstrate that the model can accurately perform high-dimensional angular integration, precisely encoding both azimuth and pitch of the bat’s head direction in 3-D space. Meanwhile, individual neurons in the HD network developed conjunctive tuning for azimuth × pitch. We recorded the firing rates of all the HD cells at each time point and decoded the head direction using Eq 17 and 18. Comparing the decoded and true head direction revealed that the HD population can accurately encode 3-D head direction in real-time using only vestibular angular velocity inputs without visual information. The directional tuning of each HD cell can be obtained by mapping the 40-minute firing activity of each individual cell to the corresponding true head direction at each time point. The results revealed that a single HD cell exhibits conjunctive encoding of both azimuth and pitch Euler angles, consistent with the tuning property of certain HD cells recorded in the bat presubiculum in 3-D space (20).

In Fig 7a, the blue trajectory represents the simulated head direction changes during the 40-minute flight. We selected three example time points (100000 ms, 200000 ms, and 500000 ms) and illustrated their corresponding azimuth and pitch values (red triangles in Figs 7b-d). The firing activity of the HD cell population at these time points is shown on the neural sheet in Figs 7e-g. The position of the activity bump on the neural sheet corresponds precisely to the azimuth and pitch coordinates in the angular space shown in Figs. 7b-d, demonstrating that the HD population activity accurately encodes the bat’s 3-D head direction (see also S1 Video for demonstration).

**Fig 7.**
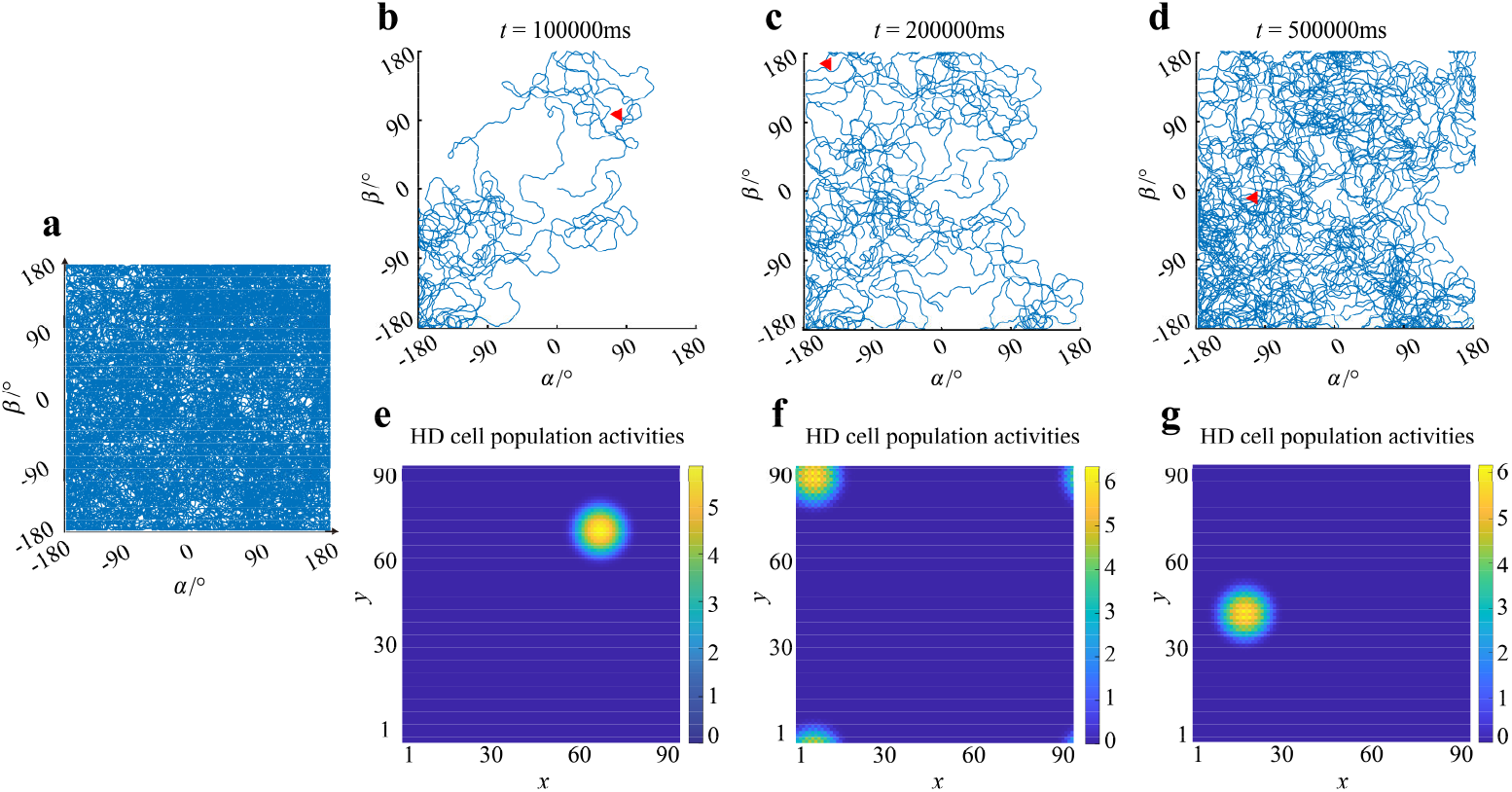
Population dynamics of HD cells tracked complex changes in azimuth and pitch of bat’s head orientation in 3-D space. (a) Trajectory of 3-D head direction changes on the angular plane (horizontal: azimuth *α*; vertical: pitch *β*). (b-d) At 100000 ms (b), 200000 ms (c) and 300000 ms (d), the head direction (red triangle) and the cumulative trajectory (blue line) of head direction trajectory up to the current time point. (e-g) Population activity of HD cells on the neural sheet at 100000 ms, 200000 ms and 500000 ms. The position of activity bump matching the bat’s head direction at each time point, demonstrating consistent accurate tracking of 3-D orientation by the model.

We further decoded the azimuth and pitch at each time point from the HD cell population activity. The decoded values were compared with the true angles to quantify decoding errors. Throughout the 40-minute simulation, the azimuth decoding error *e*_*ah*_ remained within 4° (Fig 8a, top). The decoded azimuth trajectory (orange) closely aligned with the true azimuth (blue), with almost perfect overlap as shown in the bottom panel of Fig 8a. The inset provides a magnified view of the boxed region to highlight the precision of decoding. Similarly, the pitch decoding error remained within approximately 4° (Fig 8b, top), and the decoded pitch trajectory (orange) matched the true pitch (blue) with high fidelity (Fig 8b, bottom; inset shows details). These quantitative results demonstrate that the modeled HD cell population accurately integrates high-dimensional angular velocity and encodes the bat’s 3-D head direction over long durations.

**Fig 8.**
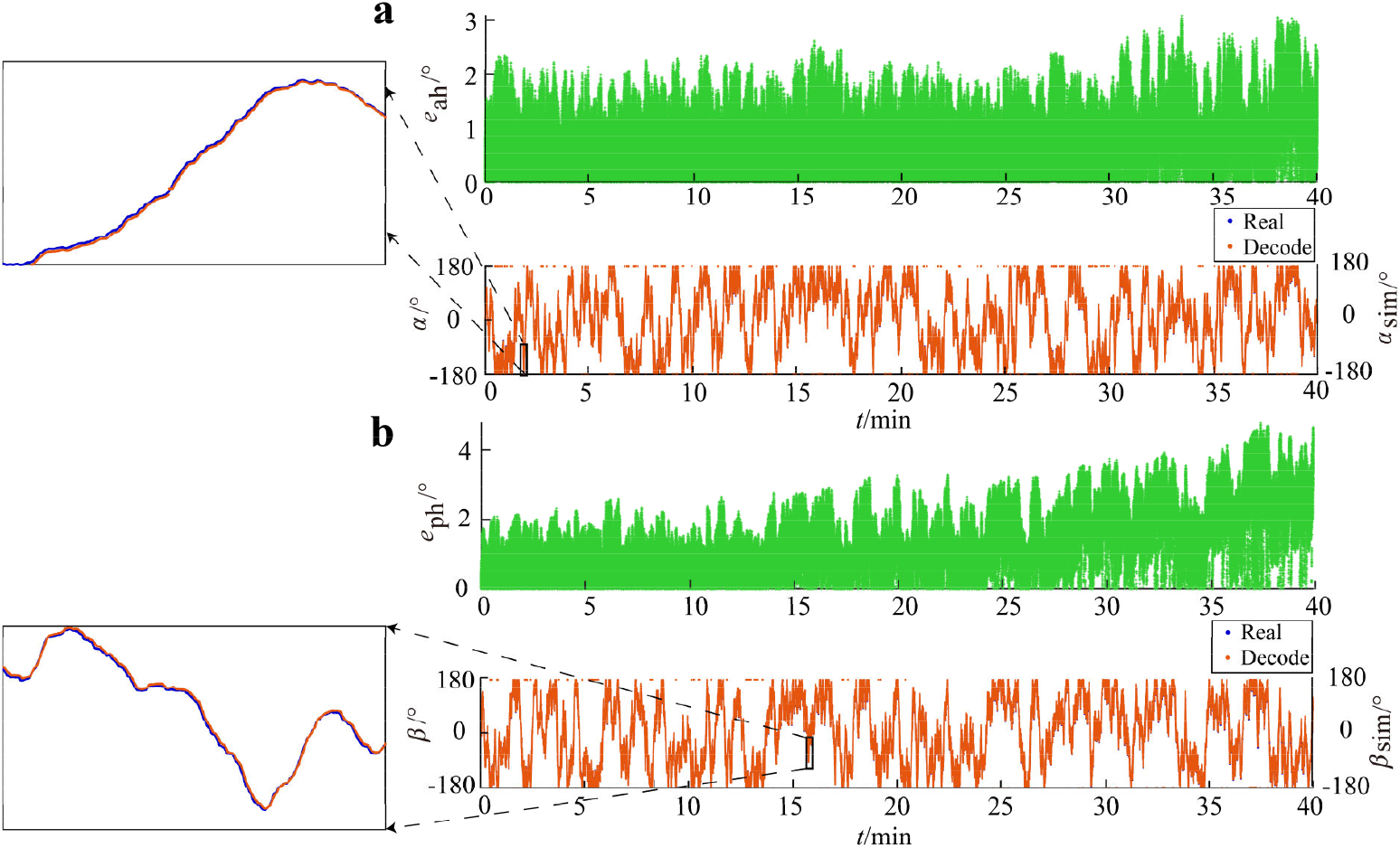
Decoding error of head directions in 3-D space by population dynamics. (a) Decoding of head direction azimuth from HD population activity during 3-D movement, showing decoded azimuth *α*_*sim*_ and decoding error *e*_*ah*_. Top: Error between decoded azimuth *α*_*sim*_ and true azimuth *α*, remaining within 3°. Bottom: Decoded azimuth (orange) versus true azimuth (blue). The trajectories overlap extensively (orange covers blue) owing to accurate encoding. The inset shows a magnified view of the region marked by the black square. (b) Decoding of head direction pitch from HD population activity, showing decoded pitch *β*_*sim*_ and decoding error *e*_*ph*_. All figure conventions share the same meanings as in (a).

### Directional tuning properties of individual HD cell in 3-D space

The 40-minute simulation of 3-D head direction changes covered the full 360°×360° angular space, enabling us to correlate the firing rates of each HD cell to all possible head directions. This yielded high-dimensional preferred direction vectors and tuning surfaces (analogous to tuning curves in lower-dimensional HD neurons) for all 8100 HD cells. Using Eqs 11-13, we computed the empirical preferred direction 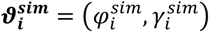 for each HD cell *i* and compared it with its pre-assigned preferred direction ***ϑ*_*i*_** = (*φ*_*i*_, *γ*_*i*_) to determine the preferred direction error 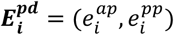. The absolute errors between the computed and preassigned preferred azimuth and pitch were within 6° for all cells. The distribution of errors for preferred azimuth across the 8100 HD cells is shown in Fig 9a (top), and the errors for preferred pitch are shown in Fig 9b (top). The bottom panels of Figs 9a and 9b provide comparisons between the pre-assigned and empirically derived preferred azimuth and pitch for all HD cells, with insets showing magnified views of the boxed regions. In fact, the close agreement between the pre-assigned and empirically derived preferred directions of individual HD cells occurs if and only if the HD system accurately tracks head direction changes. Thus, these results provide additional evidence that the model successfully performs precise angular path integration and reliably encodes 3-D head orientation.

**Fig 9.**
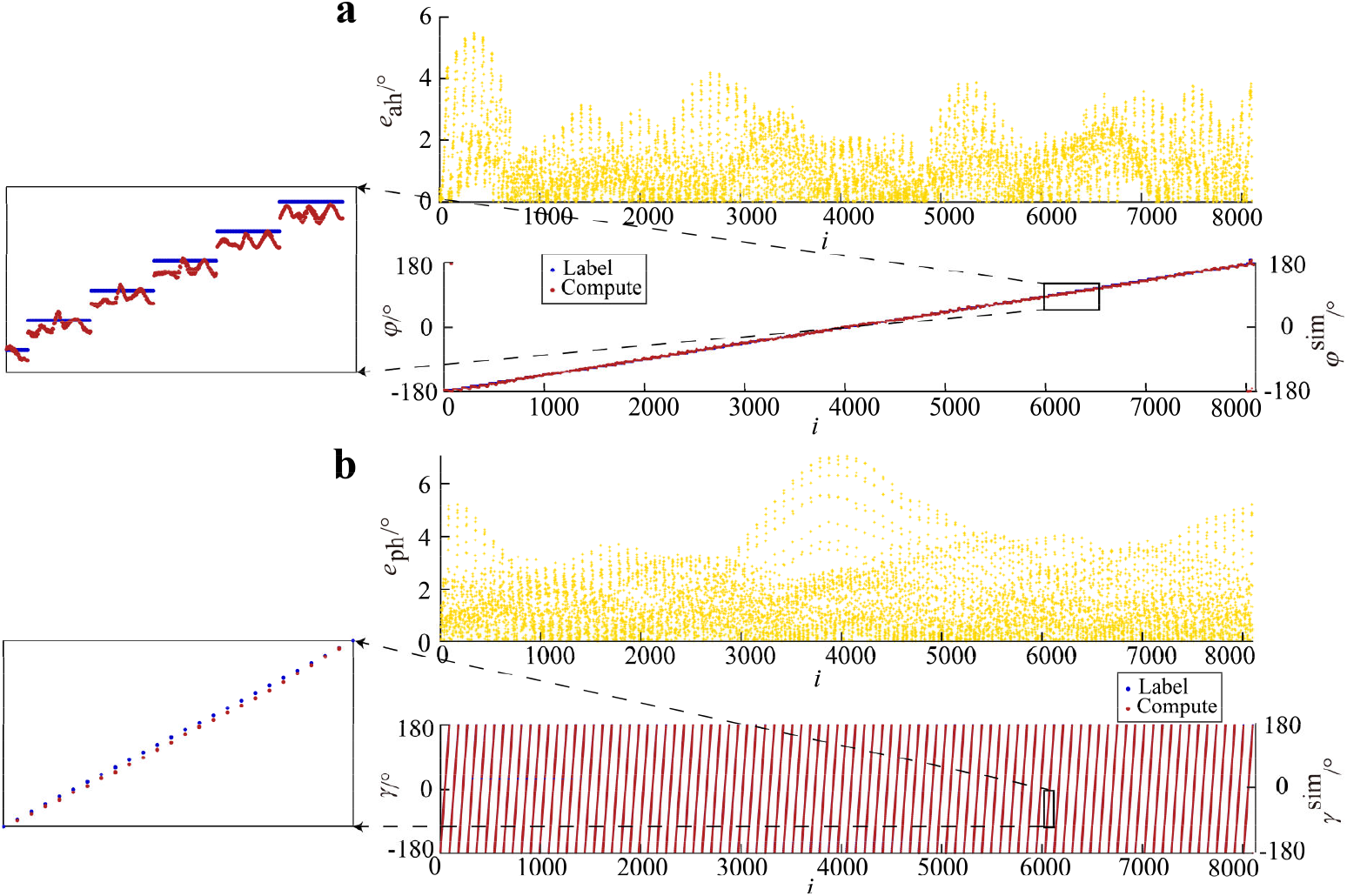
Preferred direction errors of HD cells. Based on the firing rate data of all 8100 HD cells in response to different head directions shown in Fig 7a, the empirical preferred azimuth 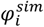 and pitch 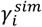 can be calculated for each cell *i*. These preferred directions were compared with their initially assigned preferred directions. (a) Computed empirical preferred azimuth 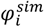 and the error 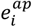 compared with preassigned azimuth for each of the 8100 HD cells. Top: Error 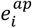 between computed preferred azimuth 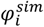 and preassigned *φ*_*i*_, limited to within 6°. Bottom: Computed 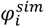 (red) versus pre-assigned *φ*_*i*_ (blue). The perfect alignment results in red points largely obscuring blue points. The inset shows a magnified view of the region marked by the black square. (b) Computed empirical preferred pitch 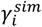 and the error 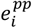 compared with pre-assigned pitch for each of the 8100 HD cells. All figure conventions share the same meanings as in (a).

The tuning surfaces of eight example HD cells are shown in Fig 10. These eight cells have pre-assigned preferred directions of (-176°, -176°), (-176°, -60°), (-176°, 60°), (-176°, 180°) and (-60°, -176°), (-60°, -60°), (-60°, 60°), (-60°, 180°) as shown in Fig 10 a-d and e-h. Their empirical preferred directions are (-176°, -176°), (-175°, -62°), (-179°, 59°), (-176°, -179°) and (-56°, -177°), (-60°, -60°), (-61°, 57°), (-56°, 179°) respectively. The tuning surfaces are shown on the upper panels of Fig 10a-h, demonstrating circular unimodal response profiles in angular space (Note that these figures illustrate time-accumulated tuning properties of individual neurons, different from previous population activity snapshots at single time points such as in Figs 7e-f). The lower panels of Figs 10a-h show the tuning surfaces mapped onto the toroidal topology, on which the azimuth and pitch coordinates are shown as red and yellow circles, respectively. The tuning surface centers for cells with preferred directions (-176°, -176°) and (-176°, -60°) show an approximately 120° difference along the pitch axis while sharing similar azimuth values (Figs 10a and b). Similarly, the centers for cells with preferred directions (-176°, -176°) and (-60°, -176°) show an approximately 120° difference along the azimuth axis while maintaining same pitch (Figs 10a and e). All HD cells in the model exhibit unimodal tuning surfaces select to 3-D head directions, consistent with electrophysiological recordings of conjunctive azimuth-pitch tuned HD cells in bats (20).

**Fig 10.**
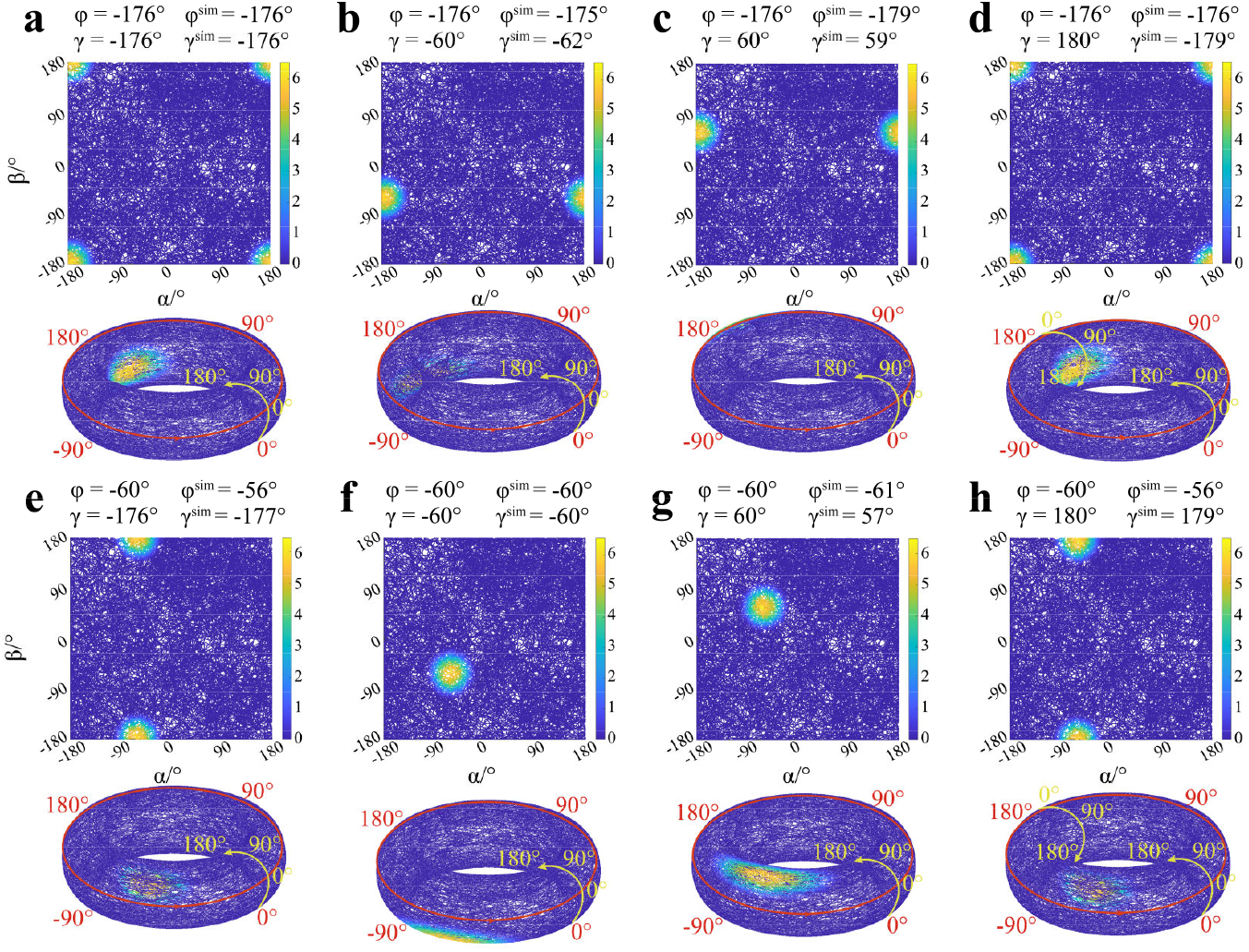
The tuning surfaces of HD cells. Tuning surfaces (analogous to tuning curves in lower-dimensional HD neurons) for eight different HD neurons. Each neuron has a pre-assigned preferred direction ***ϑ*_*i*_** = (*φ*_*i*_, *γ*_*i*_) and an empirical preferred direction 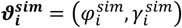 derived by correlating firing rates with head directions throughout the 40-minute simulation (Fig 7a). The upper panels illustrate the neural tuning profiles mapped onto a two-dimensional angular plane, while the lower panels show the same data onto the toroidal topology. Red and yellow circles are the circular coordinates of azimuth and pitch.

### Effects of visual cue manipulation on HD cell system in 3-D space

Experimental results from classical low-dimensional studies of HD cells show that visual information can influence the encoding of head direction (3, 6, 28-30, 49, 50). Our model confirms that the same phenomenon exists in the azimuth tuning of HD cells encoding 3-D head direction. During simulation, we initially placed the virtual visual cue at a horizontal azimuth of 0°. VIS cells encoded the visual information and transmitted it to the HD cell population through the method shown in Fig 3. After the HD cell population stabilized, it formed an activity bump on the neural sheet, thereby initializing the HD cell population. By rotating the visual cue once and multiple times in the horizontal direction, we found that the azimuth representation of HD cell population exhibited systematic errors, and the preferred azimuth of individual HD cells also shifted.

To investigate how visual cue affect the dynamics of the horizontal HD network, we simulated a 2-minute head movement trajectory. Along the horizontal direction, the azimuth changed from 0° to 90° at 0.18°/ms, and remained constant after reaching 90° (at 500 ms), while along the vertical direction, the pitch remained constant at 0° (Figs 11a-c). We placed the visual cue at a horizontal azimuth of 0° for initialization, with the head direction at (0°, 0°), and a stable activity bump formed at the center of the HD cell sheet. The visual cue was then removed, entering a dark period. The head direction system used the 0° azimuth of the initial visual cue as a reference starting point for angular integration and head direction encoding. After the head azimuth changed to 90° (*t* = 500 *ms*), the visual cue was placed at a horizontal azimuth of 90°, and visual information was again input to the HD cell layer (Fig 3e). Before the bat’s head azimuth reached 90° (Figs 11a-c), the population representation of HD cells moved consistently on the neuronal sheet to track the head direction (Figs 11d-f). When the head azimuth reached 90° and the shifted visual cue reappeared in the bat’s field of view, VIS cells encoding a horizontal angular difference of 0° provided the strongest input to HD cells whose preferred azimuth was 0°(Fig 3e). Under the influence of the mismatched visual input, the activity bump on the HD neuronal sheet gradually shifted, stabilizing again at the 0° azimuth position on the neuronal sheet at approximately 0.6 min (Fig 11i, also see S2 Video for dynamic illustration). The population representation of HD cells was shifted by about 90° under the influence of the visual cue (Figs 11g-i, Fig 12a). Equivalently, the 90° azimuth in real physical space was encoded as the 0° azimuth in the HD system.

**Fig 11.**
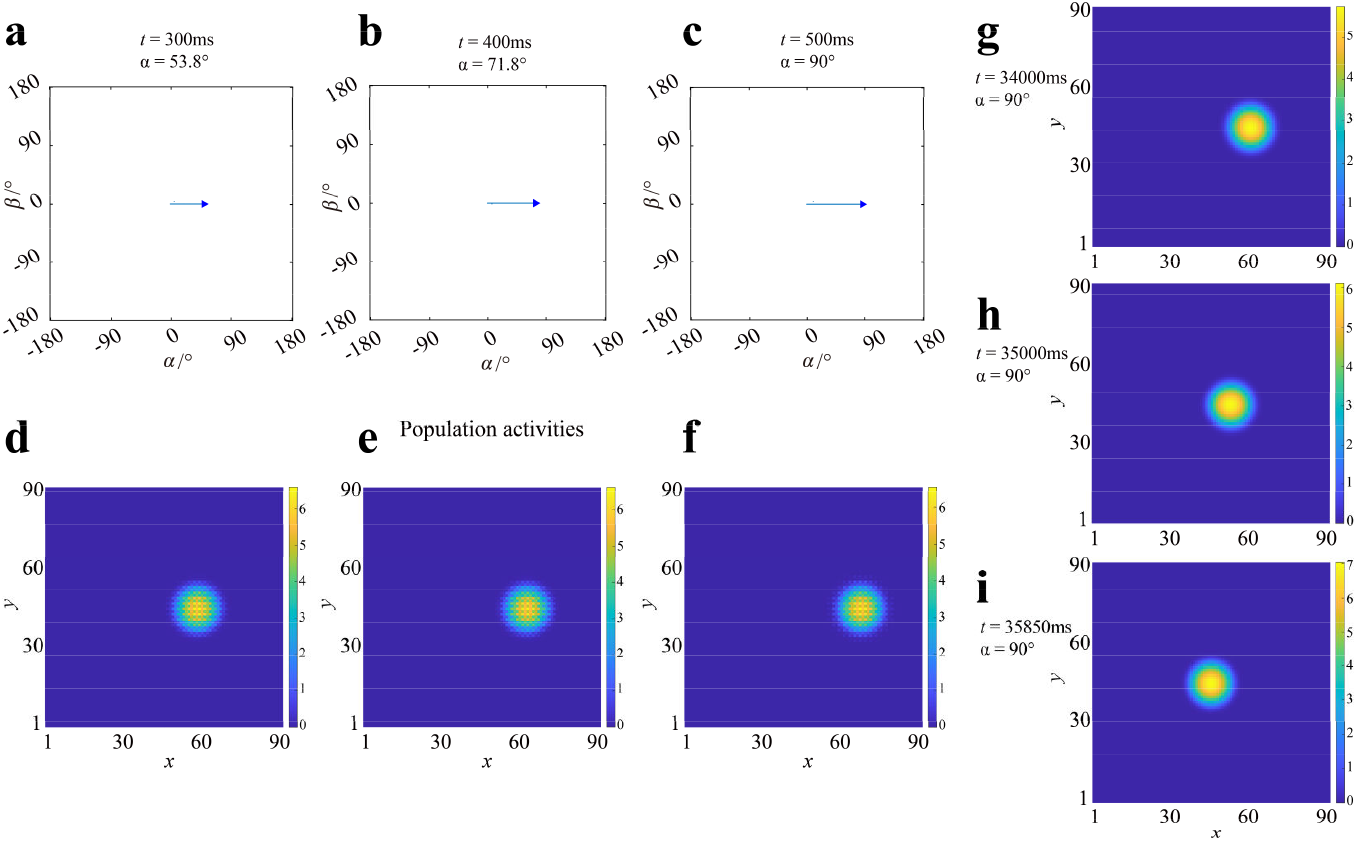
HD cells’ population representation shifted approximately 90° horizontally after visual cue rotated 90° along the horizontal direction. Along the horizontal direction, the azimuth increased uniformly from 0° at a rate of 0.18°/ms until reaching 90° (*t* = 500 *ms*) while the pitch remained constant at 0°. (a-c) Angular trajectory with changing azimuth and constant pitch (0°). Head direction trajectories at 300 ms, 400 ms, and 500 ms are shown. Blue triangle indicates the head direction at each time point, corresponding to (53.8°, 0°), (71.8°, 0°) and (90°, 0°) respectively. (d-f) Population activity of HD cells on the neural sheet at the corresponding time. The activity bump shifts consistently with the changing azimuth. (g-i) Population activity of HD cells on the neural sheet at *t* = 34, 000 *ms*, 35,000 ms and 35,850 ms, after the visual cue had been rotated to 90° azimuth at 500 *ms*. While the azimuth remains constant at 90°, the activity bump shifts back to the center of the neural sheet.

**Fig 12.**
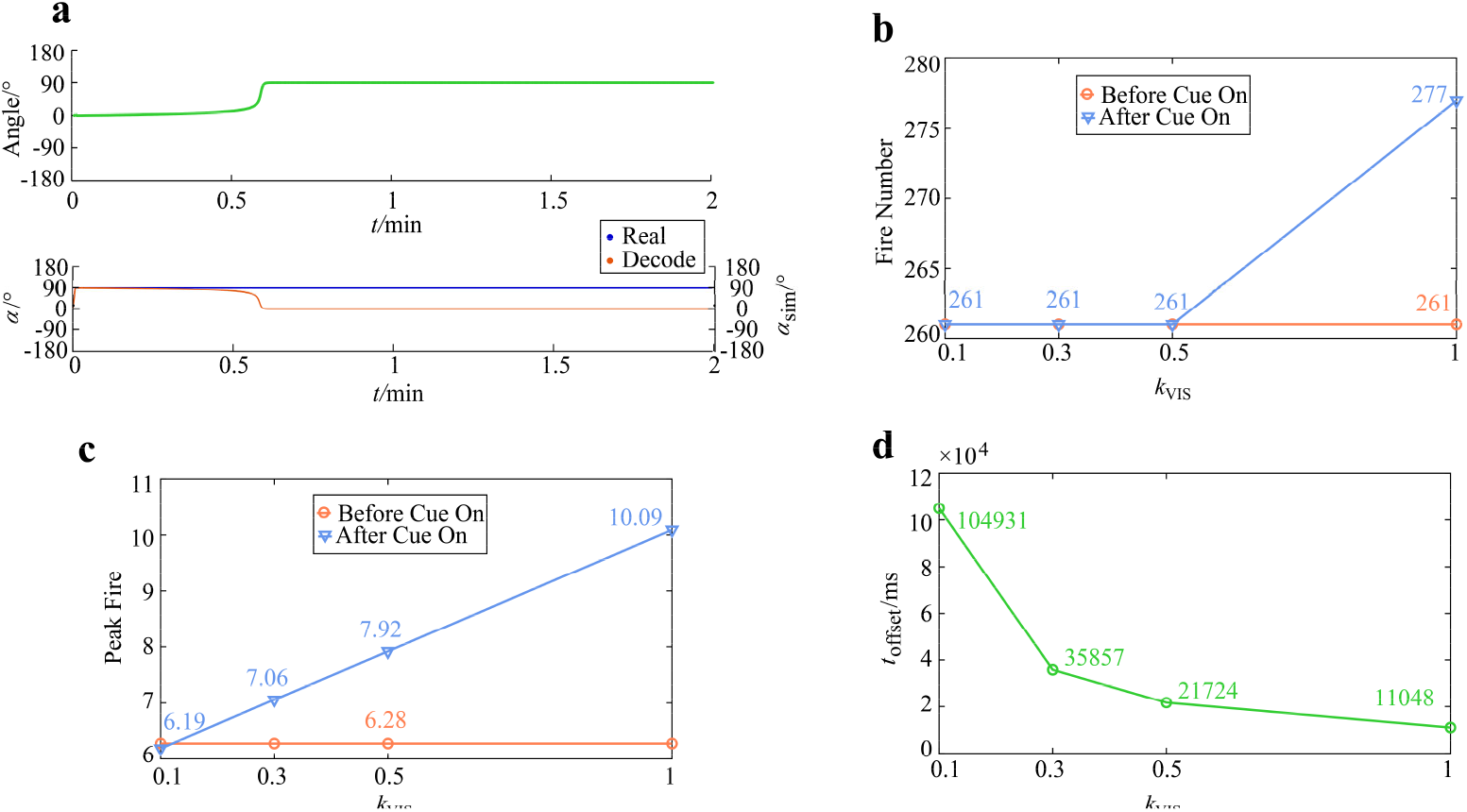
Influences of *k*_*VIS*_ on the population dynamics of HD cells. (a) Top: Comparison between the azimuth decoded from HD population activity and the true azimuth. An approximately 90° offset occurs at around 0.6 min. Bottom: Blue trajectory represents the true azimuth; orange trajectory represents the azimuth decoded from HD population activity. (b) The horizontal axis represents *k*_*VIS*_, which determines the strength of visual input to the HD cell population. The vertical axis represents the number of active cells. Orange and blue represent before and after visual cue, respectively. (c) Maximum firing rate of HD cells increases as the visual input strength *k*_*VIS*_ increases. (d) As visual cue strength *k*_*VIS*_ increases, the time required for the HD population representation to shift decreases.

To examine how visual input strength shapes HD cell population dynamics, we further simulated the model with *k*_*VIS*_ equaled to 0.1, 0.3, 0.5, and 1 (which can be regarded as an abstraction of visual salience). The results show that: 1) An increase in the strength of visual input slightly increased the number of significantly active HD cells. However, when *k*_*VIS*_ was less than 0.5, increasing its value had no effect on the number of significantly active cells (Fig 12b); 2) Visual input increased the maximum firing intensity of the HD cell population (Fig 12c), consistent with recent findings in mouse postsubiculum where objects increased firing rates of HD cells aligned with a visual object (51); 3) The strength of visual input also affected the time required for the internal representation of the HD system to produce a corresponding offset (equal to the visual cue rotation angle). As *k*_*VIS*_ increased, the time required for the HD cell population representation to develop an 90° offset became shorter (Fig 12d).

We further analyzed the effect of manipulating visual cue in the horizontal direction on head direction encoding in 3-D space using the dynamic trajectory in Fig 7a (40-minute duration). We first initialized the HD cell population by placing the visual cue at a horizontal azimuth of 0° and followed by a 5 minute dark period. Then the manipulated visual cue appeared within the bat’s field of view at an azimuth of 90°, where it remained until being removed at the 20 minute. The model then continued simulation under dark condition for additional 20 minutes. As shown in the top panels of Figs 13a and b: after 5 minute, the visual cue entered the bat’s field of view and the HD population representation shifted by 90° in azimuth (also see S3 Video for dynamic illustration). This offset persisted stably throughout the subsequent 20-minute dark period after removal of the visual cue (Fig 13a, top). Along the vertical direction, the peak decoding error for pitch increased slightly while the visual cue was present. This may be attributed to changes in the peak firing rates of activated neurons due to the visual input, which further affects the population vector, but no systematic error appeared in pitch encoding (Fig 13b, top). The bottom panels of Figs 13a and 13b show the true (blue) and decoded (orange) azimuth and pitch angles during the 40-minute simulation. A clear divergence in azimuth occurs at 5 minute, reflecting the systematic error induced by visual cue manipulation. In contrast, pitch encoding remains unaffected by the horizontal manipulation of the visual cue.

**Fig 13.**
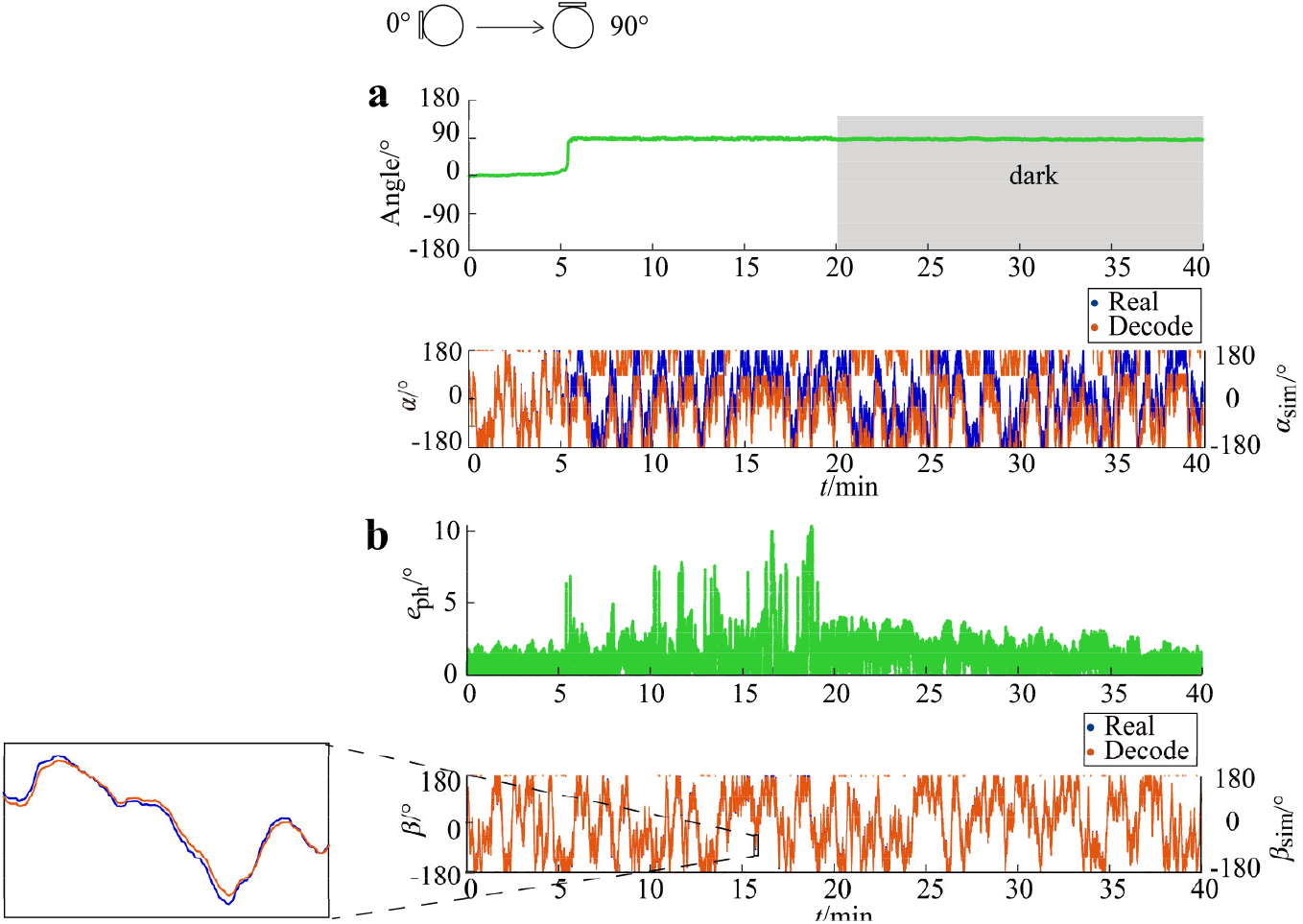
Visual cue rotated 90° along the horizontal direction in three-dimensional space, HD cells’ population representation shifted -90° accordingly. *α* and *β* represent the true azimuth and pitch of the bat’s head direction, while *α*_*sim*_ and *β*_*sim*_ represent the decoded azimuth and pitch from the activity of the HD cell population; *e*_*ph*_ indicates the decoding error for pitch along the vertical direction. (a) When the visual cue is rotated from 0° to 90°, the azimuth representation of the HD cell population shifts by 90° to the opposite direction. Top: The horizontal axis represents time and the vertical axis (Angle) indicates the offset of the azimuth representation, defined as the true azimuth minus the decoded azimuth (*α* − *α*_*sim*_), which aligns with the manipulation of the visual cue. Bottom: True (blue) and decoded (orange) azimuth. (b) During the visual input phase, the decoding error for pitch along the vertical direction slightly increases. Top: The vertical axis represents the error magnitude. Bottom: True (blue) and decoded (orange) pitch, whose values are nearly identical and the blue curve is mostly covered by the orange one. The inset shows a magnified view of the region marked by the black square. All data have been down-sampled from the original dataset for clarity of presentation.

We further investigated how manipulations of visual cue influence the directional tuning of individual HD cells. The top row of Fig 14 illustrates the tuning surfaces of HD cells with initial preferred directions of (-176°, -176°), (-176°, -60°), (-176°, 60°), and (-176°, 180°) when the visual cue is positioned at 0° azimuth. When the visual cue is rotated to 90° azimuth, the tuning surfaces obtained from activity data of the same cells shift accordingly, as shown in the bottom row of Fig 14. The peak of each cell’s tuning surface shifts along the horizontal axis by an amount matching the visual cue rotation. This result indicates that salient visual cue manipulations systematically alter the directional tuning of individual HD cells encoding 3-D head direction, a finding consistent with observations from low-dimensional rodent recordings of HD cells in postsubiculum, the anterior thalamic nucleus and other brain areas (3, 29, 50, 52).

**Fig 14.**
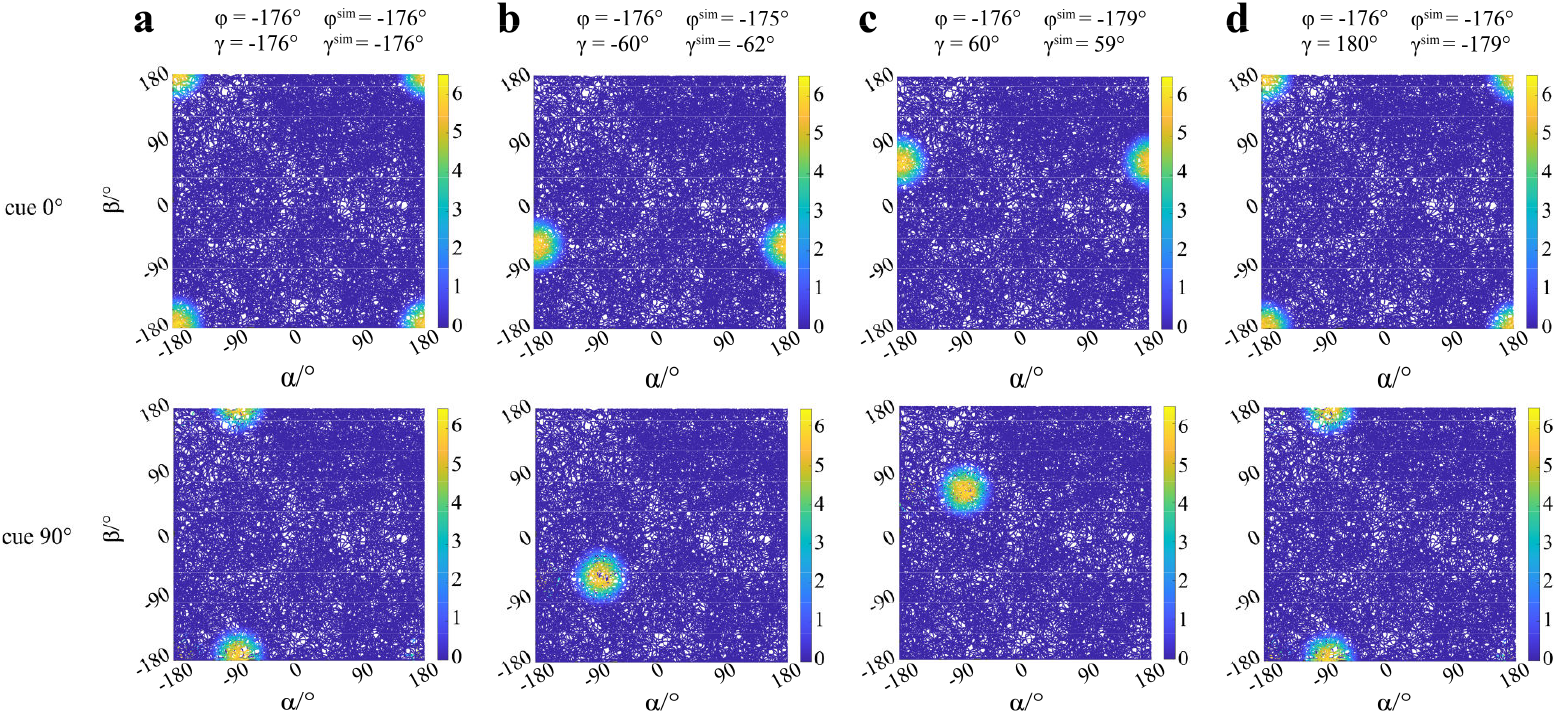
Visual cue rotated 90° along the horizontal direction in 3-D space, the preferred azimuth of HD cells shifted 90° accordingly. The tuning surfaces of four example HD neurons (same as in Fig 10) under different visual cue conditions: cue 0° (Top row) indicates the visual cue is at 0° azimuth on the horizontal plane, while cue 90° (Bottom row) indicates the visual cue is at 90° azimuth. Comparing the top and bottom rows for each column, the preferred direction in the tuning profile of each individual HD cell shifts by approximately 90°, corresponding to the rotation angle of the visual cue.

We next examined whether repeated cue manipulations in 3-D space induce cumulative errors in the encoding of azimuth by HD cell population. Using the same 40-min trajectory as in Fig 7a, we divided the simulation into four 10-min sessions. Each session consisted of an 500s light phase, during which the HD population received both visual input and angular velocity signals, followed by a 100s dark phase with vestibular input only. At the start of each session, the visual cue was rotated by 90° successively, presented sequentially at azimuths of 0°, 90°, 180°, and -90° (270°) for each session.

Decoding analysis of the population activity across all four sessions (Fig 15a, top) shows that the azimuth representation shifts systematically in accordance with the cue position, yielding offsets of approximately 0°, 90°, 180°, and -90°. These shifts persist during the dark phases until updated by a new visual cue in the next light phase. In contrast, the pitch representation remains unaffected by the horizontal cue manipulations (Fig 15b, top). The bottom panels of Figs 15a and b show the true (blue curve) and decoded (orange curve) azimuth (a) and pitch (b) during the 40-min simulation. Each cue rotation introduces a sustained discrepancy between the true and decoded azimuth, which updates with subsequent rotations. Conversely, the decoded pitch accurately tracks the true pitch throughout (Fig 15b, bottom), confirming that vertical head direction encoding is approximately independent to horizontal visual perturbations. Therefore, in 3-D space, when visual cue are repeatedly rotated horizontally, the HD cell population representation for azimuth develops successive, opposite offsets with equal magnitude as the visual cue rotations, while representation for pitch remains stable.

**Fig 15.**
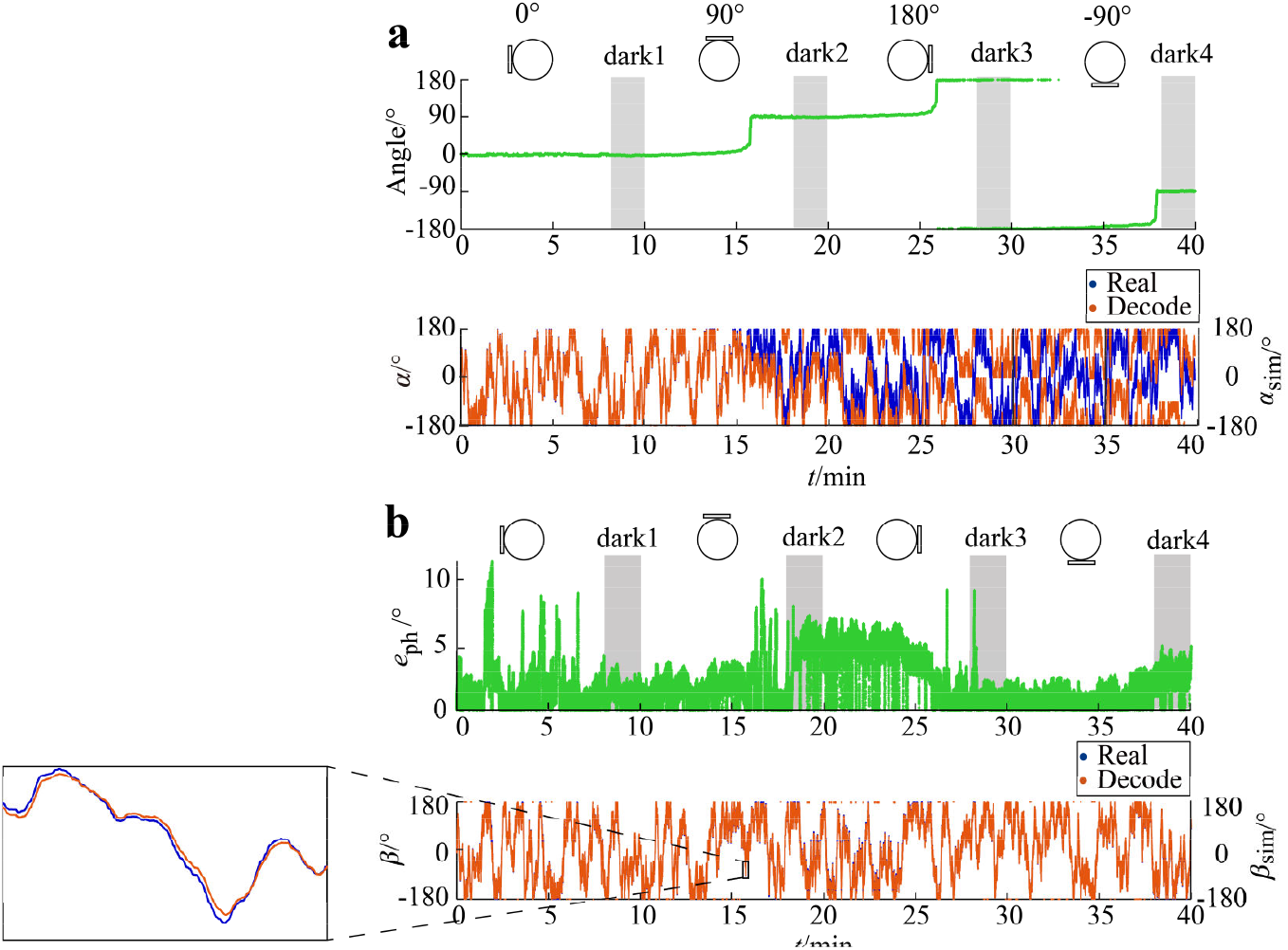
Visual cue rotated multiple times along the horizontal direction in 3-D space, the HD cells’ population representation shifted consistently. All figure features are Similar to Fig 13. *α, β, α*_*sim*_, *β*_*sim*_, *e*_*ph*_ and *angle* represent the true azimuth, true pitch, decoded azimuth, decoded pitch, pitch decoding error and azimuth offset, respectively. (a) Azimuth encoding shifts following visual cue rotations (0°→ 90°→ 180°→ - 90°). The internal HD representation develops offsets that match the cue rotation. Top: Time course of the offset (*α* − *α*_*sim*_), calculated as true azimuth minus decoded azimuth. Bottom: True (blue) and decoded (orange) azimuth. (b) Pitch encoding remains stable under azimuth visual cue manipulations. Top: Pitch decoding error fluctuates within 10° but shows no systematic shift. Bottom: True (blue) and decoded (orange) pitch. The inset shows a magnified view of the region marked by the black square. All data using for plot have been down-sampled from the original data for clarity of presentation.

## Discussion

This study presents a continuous attractor network model of HD cell in the bat presubiculum to explain their encoding of 3-D head orientation. In 2-D subspace, the model can successfully encode the horizontal azimuth, consistent with existing models of rodent HD cells (37, 48, 53-55). Moreover, the model achieves conjunctive encoding of both azimuth and pitch Euler angles in 3-D space. At the population level, the network model can accurately track simultaneous changes in azimuth and pitch across a 360°×360° angular manifold relying solely on vestibular self-motion inputs, covering most naturalistic movement patterns of bats. At the single-cell level, each model HD cell exhibits a unimodal tuning surface and selects its preferred azimuth and pitch.

We further incorporated visual inputs into the model and performed ‘virtual experiments’ in bats by manipulating visual cue, analogous to classic cue-rotation experiments in rodents(3, 29, 30, 49, 50). Simulations conducted in both 2-D subspaces and full 3-D space revealed that visual cue rotations induce systematic shifts in the HD population representation: the internal azimuth representation shifted in a direction opposite to the cue rotation by an equal magnitude, while the tuning surfaces of individual HD cells shifted in the same allocentric direction as the cue rotation by the same amount. Successive visual cue manipulations in 3-D space confirmed that under salient visual conditions, the HD system’s internal azimuth representation undergoes successive recalibrations, anchoring azimuth encoding to the visual cue. These results reproduce the cue-driven HD representation shifts observed in 2-D environments (30) and extend them to a fully 3-D framework, thus provides testable predictions.

In the dorsal presubiculum network of our model, the synaptic connection principle is that HD cells with similar preferred directions excite each other, while those with dissimilar preferences inhibit one another (48, 54). Periodic boundary conditions were imposed on the neural sheet of HD cells, describing the 360° periodicity of head direction changes for both azimuth and pitch in bats. In addition to receiving recurrent inputs from other HD cells, each HD cell also integrates angular velocity signals encoding head azimuth and pitch rotation. Recent evidence shows that mouse retrosplenial cortical neurons can track direction and speed of head turns (angular head velocity) by vestibular information (56). Although the mouse head orientation is constrained to horizontal azimuth in this research, we can anticipate that pitch angular velocity signal may also be found in mammals like rodent or bat, which will provide the higher dimensional angular velocity input required by our model. To incorporate visual information, we introduced a separate layer of VIS cells (57) that encode the difference between the azimuth of the visual cue and the azimuth of the animal’s current head direction, unlike most existing models where visual cells directly encode the allocentric direction of a visual cue (6, 49, 50). This visual information is then projected to the dorsal presubiculum HD cell population via one-to-one connections between the VIS and HD layers. This architecture can flexibly change the anchoring visual input without altering the connection from VIS cell to HD cell, enables easy and straightforward visual manipulation to test its influence on the encoding of 3-D head direction.

The visual cue appears at an allocentric azimuth of 0° and also is within the view field of the virtual animal for model initialization. VIS cells input can help to generate a single activity bump at the center of the neural sheet, indicating the HD system stabilizes at a specific position on the continuous attractor in state space, with the bump location representing the current head direction of (0°, 0°). This initialization serves as an anchoring process between the internal continuous attractor state and the external physical variable (58), and equivalently defines the pre-assigned preferred direction for each HD cell. The updating of the network’s dynamic state is jointly driven by self-motion signals and visual input but in different ways: angular velocity information from the vestibular system continuously shifts the system state along the attractor manifold for angular path integration, while visual input—particularly when in conflict with integrated self-motion cues—inhibits the existing activity bump and excites HD cells with strong VIS cell input, triggers the formation of a new activity bump, resulting in an abrupt (or a continuous but not asymptotical) update of the system state. Thus, the model predicts that salient sensory input is assigned higher confidence than angular path integration based on self-motion, enabling the (sparse) anchoring or re-anchoring of the internal dynamics of the HD system to external physical information.

Multiple types of HD cells have been recorded in the presubiculum of flying or crawling bats, which can encode all the three types of Euler angle—azimuth, pitch, and roll—either individually or in combination (20). In contrast to earlier spherical-topology models of 3-D head direction encoding (22-26) that restrict pitch to ±90°, our model employs a toroidal topology that explicitly incorporates the full angular range of pitch, making it particularly suitable for flying animals such as bats, which exhibit large-amplitude pitch maneuver during natural locomotion (45, 59). Previous models (22-26) have primarily focused on describing the encoding mechanisms of 3-D head direction in non-volumetric moving animals such as rats (13-16). These models are largely descriptive and seldom address the dynamical mechanisms underlying neuronal population activity. Furthermore, the feasibility of these models is constrained by the behavioral characteristics of rodents: when pitch exceeds 90°, most rodent HD cells lose directional tuning (10-12). This limitation is compounded by the geometric constraint known as the Berry phase effect—when the orientation of the locomotor plane deviates from the horizontal, the relationship between the directional reference frame and the world reference frame changes, introducing errors in azimuth computation (23). This phenomenon fundamentally arises from the non-commutative nature of 3-D rotations, as described mathematically by the special orthogonal group SO(3) (60).

A dual-axis model has been proposed to address the Berry phase problem, which relies on the gravitational vector (23). In such model, two axes are defined to explain the tuning property of HD cell in rat’s Anterodorsal Thalamic Nuclei (ADN): the egocentric dorsoventral (D-V) axis and the allocentric vertical axis defined by gravity. When the D-V axis rotates around the gravitational axis, the azimuth encoding is adjusted by an equal amount in the opposite direction to compensate for the possible error related to Berry phase. In addition to dual-axis model, several other computational frameworks (some are mathematically equivalent to dual-axis model) have been proposed, including the Yaw-Only (YO) model (10, 14) and the Tilted-Azimuth (TA) model (15, 22). The YO model treated vertical surfaces as extension of the floor and azimuth is computed by integrating yaw rotations only, which is sufficient to track head orientation when rodents walking on the surface of a cube (14, 24). The TA model defines azimuth on a tilted surface abide by dual-axis rule (23, 53). During 3-D navigation, the TA model decomposed orientation into 1-D azimuth and 2-D tilt, which is represented by a sphere topology, and the tilt angle has a range of only 180°. This framework is suitable for mapping head direction of rodents.

However, bats frequently exhibit a wide range of pitch head directions, varying freely within [-180°, 180°]. Compared to earlier descriptive models (22-26), the toroidal topology provides a more suitable framework for explaining the firing properties of diverse 3-D head direction cell types observed in the bat presubiculum (20). We further instantiate this toroidal topology through a continuous attractor network, focusing on providing a mechanistic explanation from a dynamical perspective for the conjunctive encoding of horizontal azimuth and vertical pitch in bats. This model also generates testable predictions and demonstrates the ability to accurately encode both azimuth and pitch during 3-D bat locomotion.

The effectiveness of the network model we propose can be interpreted at both the population and single-cell levels. At the population level, the dynamics of the HD cell ensemble accurately represent and track changes in 3-D head direction. At the single-cell level, individual HD cells exhibit stable tuning surfaces defined by preferred azimuth and pitch angles in 3-D space. The model results align with experimental observations of conjunctive azimuth-pitch tuned head direction cells in the bat brain (20). Furthermore, the model can be readily reduced to lower-dimensional configurations that encode individual Euler angles (requiring only low-dimensional integration of vestibular angular velocity signals). This scalability offers a mechanistic explanation for other types of head direction cells identified in the bat brain.

Our model further incorporates a visual cell layer that encodes the angular difference between horizontal visual cue and head direction, enabling simulation of virtual cue manipulation experiments. Simulations in 2-D subspaces reproduce findings from rodent visual cue manipulation studies (30), showing that the HD population representation shifts horizontally in a direction opposite to the visual cue rotation by an equal angular amount. In 3-D space, we performed single and successive horizontal cue rotations. The model predicts that visual manipulations induce two complementary effects: at the population level, the azimuth representation shifts opposite to the cue rotation by an equivalent angle; at the single-cell level, the individual HD cell’s tuning surface shifts in the same direction as the cue rotation by a corresponding amount. These results are consistent with experimental studies indicating that under conflict between visual cue and self-motion integration, the HD system prioritizes visual information (6, 27-30, 49, 50). Our model predicts that for head direction encoding in 3-D space, similar results may also be found in mammals such as the bat.

However, the toroidal topology applied in this model can only encode two Euler angles instead of three, thus simplifies the bat to a vector in 3-D space rather than a fully ‘articulated’ body with volume. The model accounts for changes in horizontal azimuth and vertical pitch, but does not incorporate roll. Behavioral observations indicate that during natural flight, especially in slow turn maneuvering, bats primarily adjust their azimuth and pitch, rarely perform roll motion (45, 59). Furthermore, the proportion of HD cells found to tune solely or conjunctive to roll is less than those of the other two Euler angle (20). Thus, the simplification adopted in our model is biologically acceptable. Besides, in examining how visual information influences the HD population representation, the model only incorporates manipulation of horizontal visual cue. The most salient reference for vertical orientation for all organisms on earth is gravitational vector, which is difficult to manipulate experimentally. Therefore, pitch encoding manipulation was not included in the current study, which could be an interesting topic for future research. Although a recent study claimed to disassociate visual input and gravitational signals by tilting the ground, the investigation of HD cell encoding still focused on azimuth on the tilting plane (47). Neural oscillations within the entorhinal-hippocampal circuit are crucial for rodents navigating in large environment, which support a ‘look around’ mechanism for sampling locations and modulate the spatial neurons’ encoding (61). Such phenomena have been modelled by the continuous attractor dynamic with adaptation mechanism supported by HD cells (62), which has not been include in our model. However, no experimental evidence to date has demonstrated the existence of a similar sweep mechanism (maybe more complex in 3-D) in bats, nor is it clear how such a mechanism would link HD, place and grid cells to support navigation in 3-D space. Future studies in bats should address the synergistic interactions among different spatial neurons during 3-D navigation.

In summary, we developed a continuous attractor network model for 3-D head direction encoding that achieves conjunctive representation of horizontal azimuth and vertical pitch Euler angles. At the population level, the network dynamics accurately track varying azimuth and pitch across a full 360°×360° angular space. At the single-cell level, individual HD cells exhibit well-defined tuning surfaces characterized by preferred azimuth and pitch. In investigating sensory integration mechanisms, we found that rotating horizontal visual cue induces a compensatory shift in the network’s internal representation—equal in magnitude but opposite in direction to the cue rotation. This phenomenon aligns with classical experimental observations in lower-dimensional HD systems, while also generating testable predictions for 3-D conditions. Our model provides a computational framework for understanding the neural mechanisms underlying directional encoding for bat during 3-D navigation. It offers a computational explanation of how multimodal sensory cues—specifically visual and vestibular signals—are integrated to calibrate spatial cognition, while also providing insights for the development of novel brain-inspired spatial navigation systems.

## Supporting information

S1 Video. Population activity accurately tracking head direction

S2 Video. 90 degrees visual cue manipulation (simple case)

S3 Video. 90 degrees visual cue manipulation in 3D space

